# Lipophorin receptors regulate mushroom bodies development and participate in learning, memory and sleep in flies

**DOI:** 10.1101/2021.11.10.467940

**Authors:** Francisca Rojo-Cortés, Victoria Tapia-Valladares, Nicolás Fuenzalida-Uribe, Sergio Hidalgo, Candy B. Roa, María-Constanza González-Ramírez, Carlos Oliva, Jorge M. Campusano, María-Paz Marzolo

## Abstract

**Background:** *Drosophila melanogaster* Lipophorin Receptors (LpRs), LpR1 and LpR2, mediate lipid uptake. The orthologs of these receptors in vertebrates, ApoER2 and VLDL-R, bind Reelin, a glycoprotein not present in flies. These receptors are associated with the development and function of the hippocampus and cerebral cortex, important association areas in the mammalian brain. It is currently unknown whether LpRs play similar roles in the *Drosophila* brain.

**Results:** We report that LpR-deficient flies exhibit impaired olfactory memory and sleep patterns, which seem to reflect anatomical defects found in a critical brain association area, the Mushroom Bodies (MB). Moreover, cultured MB neurons respond to mammalian Reelin by increasing the complexity of their neurites. This effect depends on LpRs and Dab, the *Drosophila* ortholog of the reelin signaling adaptor protein Dab1. In vitro, two of the long isoforms of LpRs allow the internalization of Reelin.

**Conclusions:** These findings demonstrate that LpRs contribute to MB development and function, supporting the existence of LpR-dependent signaling in *Drosophila*.

## Background

The Low-Density Lipoprotein Receptor (LDL-R) family is an ancient protein family conserved throughout evolution [1, 2]. The LDL-R family is widely known because of its role in lipoprotein endocytosis, controlling the systemic homeostasis of cholesterol by binding to Apolipoprotein E (ApoE) and apolipoprotein B (ApoB) [3, 4]. In insects, lipid uptake is mediated by Lipophorin Receptors (LpRs), membrane proteins belonging to the LDL-R family that are internalized and recycled [2, 5]. Lipophorins are the invertebrate’s molecular entity equivalent to lipoproteins in vertebrates and bind LpRs through their protein component (apolipophorin) [5].

At least two LpRs, LpR1 and LpR2, have been described in the genome of *Drosophila melanogaster* [6, 7]. These receptors have been linked to lipid uptake in the context of oogenesis and neuronal activity, which is consistent with the known roles of several LDL-R family members in vertebrates [6,8–10].

Structurally, LpRs consist of seven to eight LA modules, three EGF modules, a β-propeller, an O-glycosylation site near the transmembrane domain, and a cytoplasmic tail [6]. Six isoforms for LpR1 and five for LpR2 have been described in *Drosophila*. These are generated by alternative use of two distinct promoters, proximal or distal [6]. The main consequence of the use of the proximal promoter is that the resulting transcript lacks exons 1 to 3, which encode for a non-conserved N-terminal region and the first LA-module [6]. Thus, the isoforms transcribed from the proximal promoter are also known as the “short” ones, while those transcribed from the distal promoter are known as the “long” isoforms [6,9,11].

Only a subset of LpRs isoforms mediate lipid uptake [6,9–11]. Several reports link LpRs to functions beyond lipid metabolism. The first one showed that *Drosophila* LpR1 increases the immune response by modulating the uptake and degradation of Necrotic Serpin, a protein that controls the innate response to gram-negative infection in insects [7]. A second study showed that LpRs play a role in dendrite morphogenesis in larval ventral Lateral Neurons (LNv), a neuronal group that is part of the circuit that controls circadian rhythms in *Drosophila* [12]. Moreover, in the case of LpR1, this effect depends on the short isoforms. Specifically, it was reported that knocking down a specific LpR1 short isoform (LpR1G) or LpR2 in LNv reduces activity-dependent dendrite arborization [11, 12]. Since a genome-wide RNAi screen in cultured neurons from *D. melanogaster* embryos identified LpR2 as a gene implicated in neuronal development [13] and given the fact that LpR1 and LpR2 are widely expressed in the *Drosophila* larval brain [11, 12], it is possible to propose that LpRs could contribute to the development, maturation and operation of several fly brain regions.

*Drosophila* LpR1 and LpR2 share with vertebrates LDL-R, VLDL-R, and ApoER2, motifs involved in intracellular trafficking and signaling, like the NPxY [2, 6]. In this regard, besides their role as lipoprotein receptors, VLDL-R and ApoER2 serve as receptors for the secreted glycoprotein Reelin, an extracellular ligand with essential roles in the central nervous system (CNS) [14–17]. Reelin functions include neuronal polarization and migration in critical association areas in the brain, such as the cerebral cortex, hippocampus, and cerebellum. In adult animals, Reelin is also implicated in synaptic plasticity, learning, and memory [15,16,18–24].

Here, we decided to advance on the characterization of the expression and function of LpRs in the fly brain. First, we showed that *Drosophila* mutants for LpR1 and LpR2 exhibit olfactory memory and sleep architecture deficits. Since these behaviors depend on a crucial associative region in the fly brain, the Mushroom Bodies (MB), we assessed whether LpRs expression has any consequence on the structure of this fly brain area. Our data show relevant alterations in the MB organization of flies mutants for LpRs and also of animals where an RNAi for these receptors is directed to this brain region. Further, the alterations found in flies mutant for LpR1 were prevented in MB expressing LpR1J, one of the long isoforms of the receptor. Surprisingly, *Drosophila* MB neurons in primary culture were responsive to vertebrate Reelin by increasing their neurite tree’s complexity, as vertebrate neurons do. This effect depends on LpR1 and LpR2. Also, we show that cells bearing long isoforms of LpR1 and LpR2 internalize Reelin. In line with the idea of a Reelin-induced LpR-dependent response in flies, we described that MB organization and function, and the response of cultured MB neurons to Reelin, are affected by the absence of the cytoplasmic protein Dab, the fly homolog of Disabled 1 (Dab1) [25–27], which is the first component of the Reelin intracellular signaling cascade [28]. Overall, our results show, for the first time, that LpR1 and LpR2 contribute to *Drosophila* MB organization and function. Besides, our data suggest that both receptors would participate in a novel signaling cascade involving Dab, which can be activated in culture by vertebrate Reelin.

## Results

### Fly mutants for Lipophorin receptors exhibit impaired behaviors associated with MB function

In order to characterize the role of LpRs in the *Drosophila* brain, we first asked whether mutants for these receptors exhibit any alteration in one of the most studied behaviors in flies, aversive olfactory memory. We used a mutant for each LpR, generated by deleting a segment of the encoding genes [6]. These tools were validated through qPCR (S1 Fig A-B). We carried out a protocol to generate mid-term aversive olfactory memories, as explained in the Methods section (Fig 1A). Briefly, flies were exposed to two different odorants sequentially, but only the first odorant was paired to electric shocks. One hour after finishing the training protocol, the flies were allowed to choose between the two odorants. In control animals, the olfactory memory measured as performance index reached a magnitude of 0.44 ± 0.04, consistent with the idea that the training protocol generates new olfactory memories. Remarkably, the performance index recorded in the LpR1 and LpR2 mutants suggests that no memories were generated in these flies (0.00 ± 0.07 and 0.07 ± 0.14, respectively; p<0.01 as compared to controls; Fig 1B). Additionally, as experimental controls, we evaluated the mutants’ ability to respond to the conditioning stimulus (electric shocks; S2 Fig A) and the odorants used (S2 Fig B and C). Interestingly, LpR1 mutants display a higher olfactory acuity towards one of the odorants used (S2 Fig B), which is one of the reasons we carried out the training protocol using either odorant as a conditioned stimulus for different experiments.

**Fig 1.**
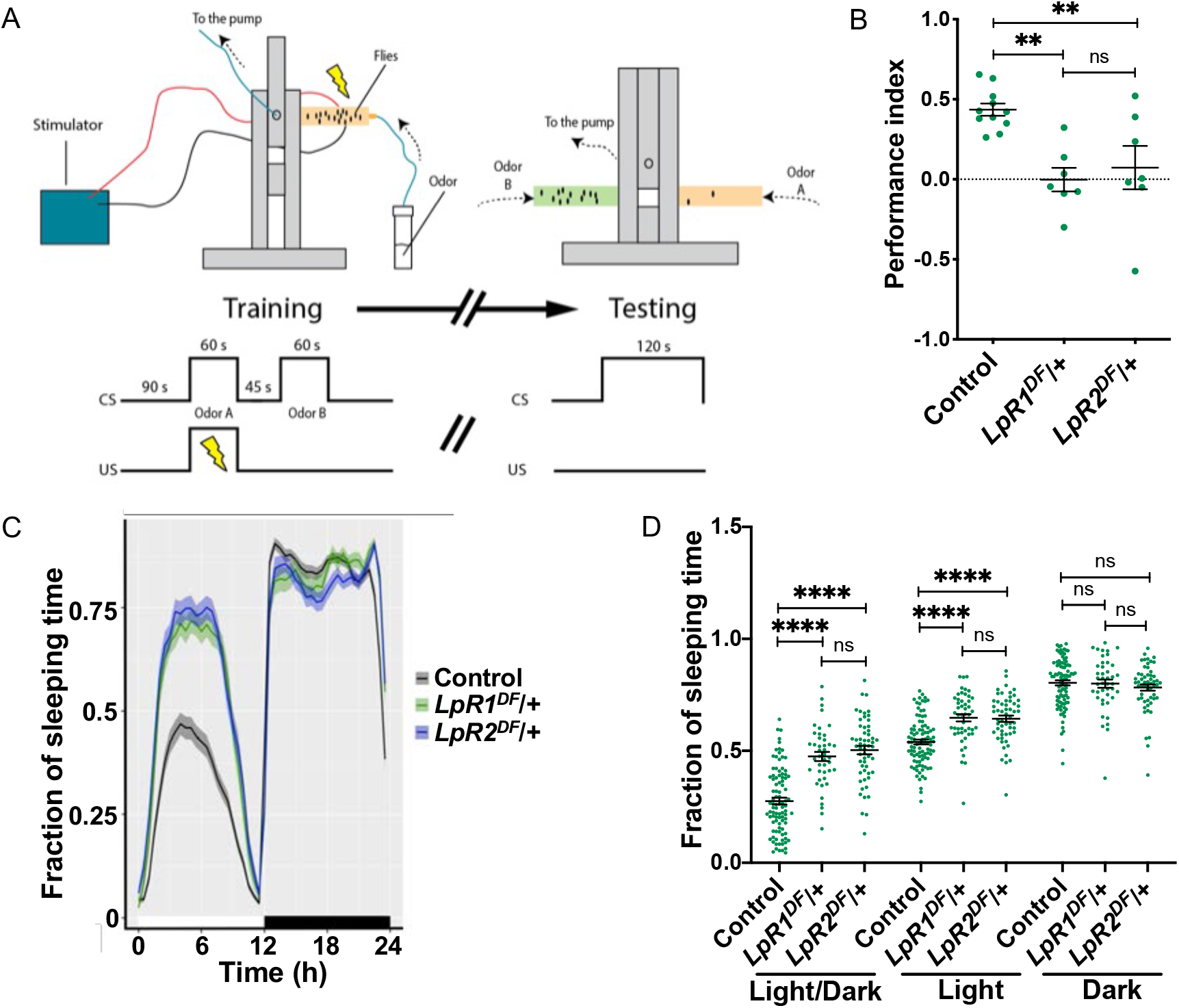
Impairment in MB-associated behaviors in LpR1 and LpR2 mutants. **A** Scheme of the setup and protocol for aversive olfactory memory assay. Flies were allowed to familiarize themselves with the training apparatus for 90 s. Then, flies were exposed to an odorant (conditioned stimulus, CS+) over 60 s, while they received electric shocks (unconditioned stimulus, US) (left panel). After this, flies were allowed to rest for 45 s in the presence of fresh air. Afterward, flies were exposed to a different odorant for 60 s, which was not paired to electric shocks. One hour after this training, memory was evaluated. Memory test (right panel) consists of having the flies in the center of the apparatus where they are exposed to the two odorants (each in one arm) for 120 s. In that period, flies are allowed to choose between them. **B** Performance index for olfactory memory in control animals and mutant flies for LpR1 (*LpR1^DF^*/+) or LpR2 (*LpR2^DF^*/+). n= seven experiments carried out in independent groups of flies. One-way ANOVA followed by Dunnett’s test, p=0.0011. ** indicates p<0.01, “ns” not significant. **C** Sleep profile throughout the day in flies mutant for LpR1 or LpR2. Above the X-axis, the white bar represents the hours of the day when flies were exposed to light; the black bar represents the hours when flies were exposed to darkness. **D** Fraction of sleeping time in flies mutants for LpR1 and LpR2. Two-way ANOVA shows that light condition and genotype factors, and also interaction between these factors, play a role in results (p<0.0001 for each analysis); Tukey post-test; *, ** and ****, indicates p<0.05, p<0.01 and p<0.0001 between conditions. ns, not significant. Data in **C** and **D**, from two independent experiments, were from n=43-95 flies studied per genotype. In **B**-**D**, data expressed as mean ± SEM.

We then assessed circadian rhythms in LpRs mutants while under a light-dark cycle of 12-12 hours [29, 30]. Interestingly, the observation of actograms supports the idea that flies mutant for LpR1 and LpR2 exhibit increased sleeping time than control flies, particularly during the light phase (Fig 1C-D). A summary of data collected over six days evidenced higher sleep time in both mutant strains during the light phase (Fig 1D).

Altogether, these results support the idea that tampering with the expression of LpRs has functional consequences reflected by impairment in memory formation and the regulation of sleep patterns.

### Lipophorin receptors contribute to adult Mushroom Body organization

The MB is a well-characterized *Drosophila* structure located in each side of the brain, linked to crucial associative fly behaviors, including olfactory learning and memory, sleep, and locomotion [31–35]. Therefore, we decided to study whether alterations in olfactory memory and sleep homeostasis in mutants for LpRs are associated with changes in MB anatomy.

The principal neurons in the MB, the Kenyon cells, organize their dendrites and axons to give rise to the Calyx, peduncles, and lobes (Fig 2A) [36, 37]. The Calyx, localized in the posterior aspect of the fly brain, is a structure formed by the Kenyon cells soma and dendrites, while their axons project frontally, arranged in a bundle called the peduncle. Before the end, the axonal fibers turn perpendicularly to give rise to the so-called MB lobes. The lobes are classified in γ, α/β and α’/β’, where β, β’ and γ lobes are positioned horizontally while α and α’ lobes are placed vertically [38]. This anatomical organization has functional roles in several behaviors [39–41], so we paid close attention to MB organization in our studies.

**Fig 2.**
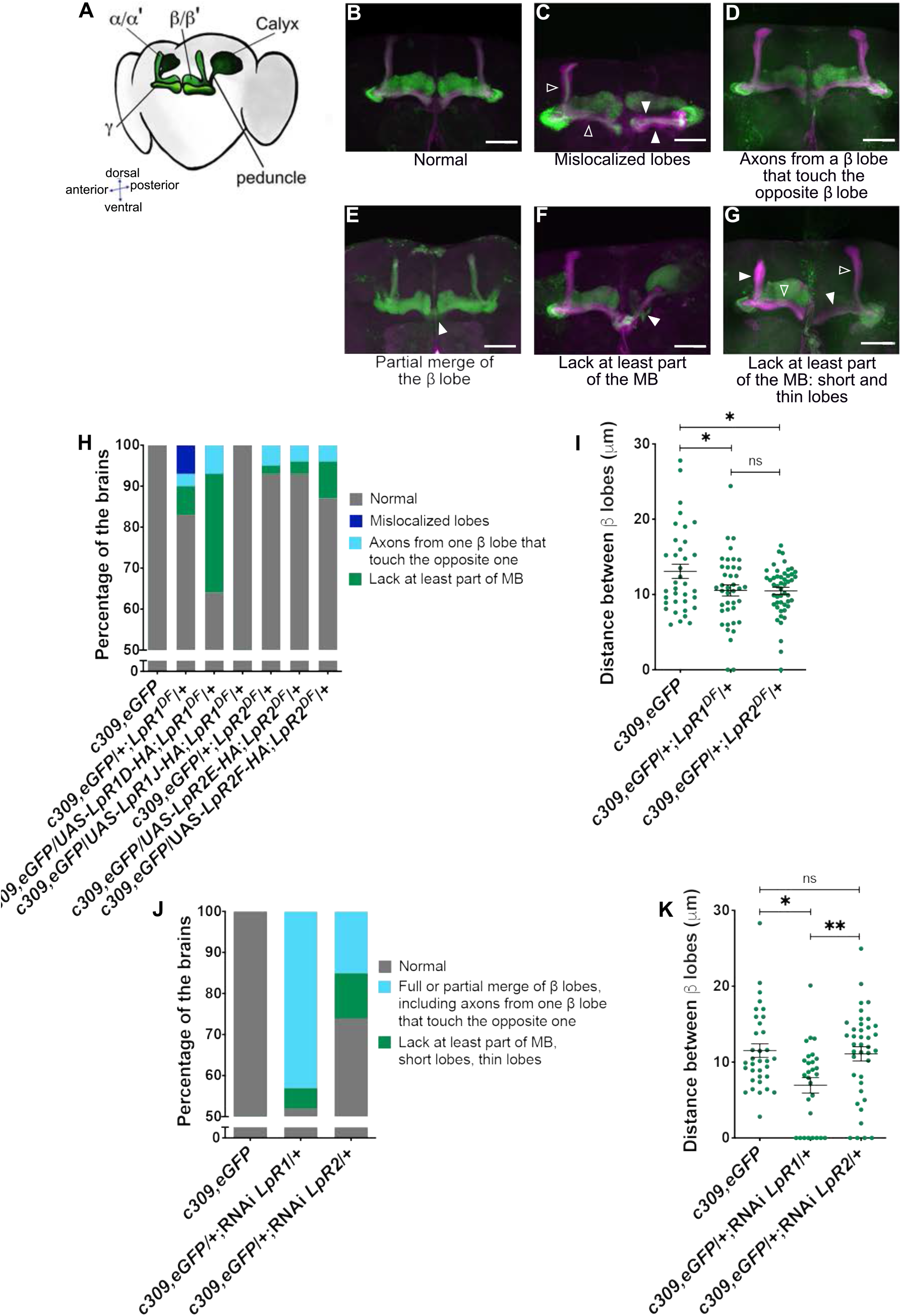
Knocking down LpR1 or LpR2 results in MB anatomical defects. **A** Scheme of the normal organization of the MB in the adult fly brain. MB subregions are identified. **B**-**G** Representative images of adult *Drosophila* brains in LpR1 or LpR2 hypomorphic mutants; in green is shown GFP expression in MB neurons (*c309,eGFP*), in magenta is presented FasII staining to visualize the MB structure. Except **F**, all microphotographs correspond to a limited number of optical sections that help observe the phenotypes identified under each image. Empty arrowheads point to normal structure in MB lobes, while full arrowheads show alterations. Scale bar: 20 μm. **H** Percentage of fly brains from each genotype exhibiting the identified phenotypes, and reversion of phenotypes by LpR expression recovery in MB. Data obtained from *c309,eGFP* control animals in n=40 brains from 4 independent experiments; *c309,eGFP*/+;*LpR1^DF^*/+ n=41 brains, 4 experiments; *c309,eGFP*/UAS-LpR1D-HA;*LpR1^DF^*/+ n=28 brains, 4 experiments; *c309,eGFP*/UAS-LpR1JHA;*LpR1^DF^*/+ n=19 brains, 3 experiments; *c309,eGFP*/+;*LpR2^DF^*/+ n=46 brains, 4 experiments, *c309,eGFP*/UAS-LpR2E-HA;*LpR2^DF^*/+ n=28 brains, 3 experiments, and *c309,eGFP*/UASLpR2F-HA;*LpR2^DF^*/+ n=23 brains, 3 experiments. Fisher test for *c309,eGFP, c309,eGFP*/+;*LpR1^DF^*/+, *c309,eGFP*/UAS-LpR1D-HA;*LpR1^DF^*/+, *c309,eGFP*/UAS-LpR1JHA;*LpR1^DF^*/+, P=2.2*10^-16^. Fisher test for *c309,eGFP, c309,eGFP*/+;*LpR2^DF^*/+,*c309,eGFP*/UAS-LpR2E-HA, *c309,eGFP*/UAS-LpR2F-HA;*LpR2^DF^*/+, P=0.0036. **I** Distance between β lobes in each side of the fly brain in the different genotypes. Data expressed as mean ± SEM from the number of brains identified in **H**; one-way ANOVA, Kruskal Wallis post-test, p=0.0028. * and **, p<0.05 and p<0.01 respectively; “ns”, not significant. **J** Percentage of brains exhibiting phenotypes in flies expressing RNAi for LpR1 or LpR2 in MB neurons. Fisher test, P=2.2*10^-16^ **K** Distance between β lobes in knockdown strains. Data in **J** and **K** from *c309,eGFP* control flies n=26 brains, 4 independent experiments; *c309,eGFP*/+;RNAi-*LpR1*/+ n=21 brains, 4 experiments; and *c309,eGFP*/+;RNAi-*LpR2*/+ n=27 brains, 4 experiments. One-way ANOVA, followed by Kruskal Wallis test p=0.0028; * and **, p<0.05 and p<0.01 respectively; “ns”, not significant.

We evaluated fly brains from adult animals deficient in LpR1 or LpR2 expression. Our strategy comprised two different genetic tools for each gene: a mutant for each LpR (described previously in the functional experiments), and RNAi expression against the receptors’ transcripts, specifically in the MB, using the Gal4-UAS system. The RNAis were validated by qPCR using the pan-neuronal driver Elav (S1 Fig C-D).

In the analysis of results obtained in mutants for LpR1 or LpR2, we found and classified a variety of phenotypes (Fig 2C-G) as compared to control brains (Fig 2B). These include mislocalized lobes (Fig 2C) in which the positions of one or more lobes are altered; the classification includes guidance defects where the α or β axons are misguided along β or α axons, respectively, resulting in thicker lobes. In some cases, we observed that the distance between MB β lobes is reduced (Fig 2D), while in other brains, we detected fascicles of axons from indistinguishable origin crossing the middle line, which results in partial or total β lobes fusion (Fig 2E). Further, we detected brains in which a part of the structure of the MB was lost; for instance, the absence of the lobes (Fig 2F), the presence of thinner or shorter lobes (Fig 2G), and very rarely, the complete lack of the MB (data not shown). These phenotypes were classified, as shown in Fig 2H-I and S1 Table.

In general, flies lacking one copy of the *LpR1* gene (*c309,eGFP*/+; *LpR1^DF^*/+) showed a higher percentage (17%) of brains exhibiting any alteration in MB than flies lacking one copy of *LpR2* (*c309,eGFP*/+;*LpR2^DF^*/+): (6.5 %) (Fig 2H). The analysis of the distance between β lobes indicated that this defect is present in mutants for LpR1 and LpR2 (Fig 2I). To further address the importance of LpR1 and LpR2 specifically expressed in the MB, we evaluated potential defects in this structure in adult flies expressing RNAi under the control of the c309 driver (Fig 2J, K and S2 Table). Similar to data obtained with LpRs mutants, the percentage of brains with MB alterations was higher in animals LpR1 knockdown (around 48 %) as compared with LpR2 knockdowns (around 26 %) (Fig 2J). On the other hand, the distance between MB β lobes was found significantly diminished only in animals expressing the RNAi for LpR1 (Fig 2K).

To confirm that these defects were triggered by the reduced expression of LpRs and not by an artifact of the MB driver used (*c309*-Gal4), we expressed the RNAi for the LpRs under the control of a second driver: *OK107*-Gal4 [42] (S3 Fig and S3 Table). This driver is strongly expressed in the whole MB but also exhibits significant expression in some neurons that do not belong to the MB [36]. The structure of MB was similarly affected in animals expressing RNAi for LpR1 or LpR2 (phenotypes were observed in 50% and 41.4% of the brains analyzed, respectively) (S3 Fig A). It was evident that the OK107 driver was associated with more severe phenotypes, including the disappearance of entire lobes. Similar to what was found using the c309 driver, the distance between β lobes was reduced in the MB after LpR1 knockdown (S3 Fig B).

In order to evaluate whether the phenotypes observed rely on the lack of specific isoforms of LpRs, we re-expressed in MB one of the short or the long versions of LpR1 (LpR1D or LpR1J, respectively) in the deficient genetic background. Besides, in the LpR2 deficient animals, we re-expressed one of the short or long isoforms of LpR2 (LpR2F or LpR2E, respectively) (for a schematic representation of receptors, see S4 Fig). For doing this, we used the Gal4-UAS system and the c309 driver. Only the LpR1J (long isoform) was able to rescue the morphologic alterations of LpR1 mutant animals. On the contrary, neither LpR2E nor LpR2F rescued the mutant phenotype in the LpR2 deficient background (Fig 2H). Overall, these results support a role of LpR in the anatomy of the MB, specifically the relevance of LpR1 and probably the requirement of an LpR2 isoform distinct from LpR2E and F.

### Lipophorin receptors contribute to Mushroom Body development and organization in the adult

Our results support the idea that LpRs contribute to the establishment of the typical anatomical organization of adult *Drosophila* MB so that flies deficient in these receptors exhibit alterations in this brain structure. To advance on this idea, we decided to study the expression of LpR1 and LpR2 in MB at distinct developmental stages in *Drosophila*. For performing this task, we used some of the tools described above.

By using the Gal4-UAS system to command the expression of fluorescent proteins under the control of an LpR1 driver, we found that in the adult brain, LpR1 was highly enriched in a group of neurons found adjacent to the MB β/β’ lobes and in the central complex (Fig 3A, inset), while its expression in the rest of the brain was rather diffuse. The cells showing the highest LpR1 expression in the fly brain seem to be the neurons projecting to the ring of the ellipsoid body [43–45]. As an alternative approach, we used a LpR1 antibody previously reported [6] and FasII staining to identify the MB (S5 Fig). Both strategies provided similar results. We also assessed the expression of LpR2 in the adult fly brain, employing a protein trap line generated by Recombination-Mediated Cassette Exchange (RMCE). In this line, the RMCE cassette is placed between the exons 12 and 13 of LpR2 and contains a GFP coding-exon that tags all the LpR2 isoforms [46]. Our data showed that LpR2 was not enriched in any specific brain region (Fig 3B). These results suggest that both LpR1 and LpR2 exhibit a rather diffuse expression throughout the adult fly brain, although LpR1 is highly expressed in a group of cells surrounding the MB.

**Fig 3.**
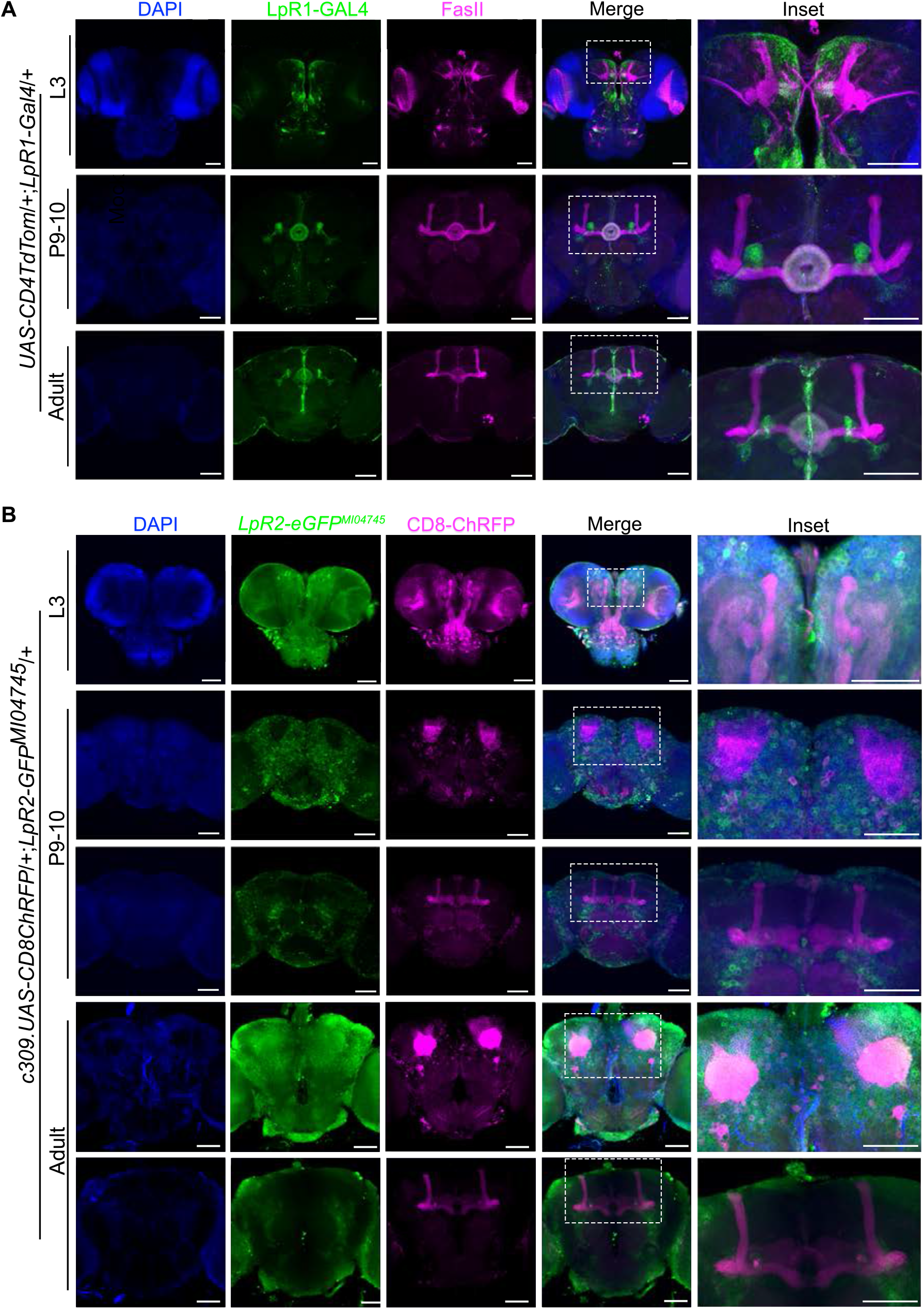
LpRs expression in the brain throughout development. **A** Representative image of a fly brain that expresses *CD4::tdTomato* (green in the image) under the LpR1-Gal4 driver in third instar larvae (n=11 brains), pupae stage 9-10 (n=2 brains) and adult animals 1-5 days old (n=10 brains). Immunofluorescence against FasII is shown in magenta and DAPI staining in blue. A discontinuous white line rectangle indicates the zone magnified in the insets. Scale white bars represent 50 μm. **B** Representative image of brains from third instar larvae (n=16 brains), pupae stage 9-10 (n=15 brains) and male adult animals 1-5 days old (n=13 brains) that expresses *LpR2-GFP^MI04745^* (green in the image) and CD8-ChRFP in the MB under the control c309 driver (magenta in the image). DAPI staining is in blue. Discontinuous white line rectangles indicate the zone magnified in the insets. Scale white bars represent 50 μm.

Considering the evidence obtained in adult flies, we analyzed the expression of LpRs in MB over larval and pupal stages, posing the idea that the alterations in adult MB structure could be explained by a modification in the developmental program that gives rise to this fly brain region. The MB begins to develop in the embryonic stage, starting from four neuroblasts per hemisphere that divide until the late pupa stage. The γ lobe develops until the mid-third instar larval stage. The α’ and β’ lobes are generated by the end of the third larval stage, while the development of α and β lobes reach maturity at the pupal stage [47, 48].

Using the LpR1 Gal4 tool, we assessed the expression of LpR1 in the larval and pupal stages. In the larval stage, LpR1 was found at low levels in the β’ lobe, with a widespread expression in the midline area and the photoreceptors, while in pupae, the receptor was found highly expressed in the central complex and in a group of neurons that surround the β’/β lobes (Fig 3A), similarly to what was found in adult flies. On the other hand, the RMCE tool was used to assess the expression of LpR2 in the larval and pupal stages. In larvae, LpR2 was observed in cells located in the midline area and the photoreceptors, whereas in pupae, it was observed in cells in the brain cortex (Fig 3B). These results indicate that LpR1 and LpR2 expression is widely distributed in the fly brain cells throughout development.

Taken together, these results suggest that LpR1 and LpR2 play a relevant role in the proper development and establishment of the adult brain structure.

### LpRs contribute to *Drosophila* MB neurite development

The proposed orthologs for LpR1 and LpR2 in vertebrates are ApoER2 and VLDL-R [2, 6]. These membrane proteins are the primary receptors for Reelin in CNS [16,22,49] and participate in the development and function of the cerebral cortex and hippocampus, among other brain areas. Besides LpRs, *Drosophila* has orthologs for other proteins involved in Reelin signaling, including the adaptor protein Dab (see below). Thus, we proposed that *Drosophila* neurons in culture would respond to mammalian Reelin, increasing their neurite complexity, as reported in cultured rodent neurons [50–53]. Strikingly, we found that our prediction was correct: MB neurons significantly increased their neurite branching in response to mammalian Reelin compared to effects induced by mock-media (Fig 4). Moreover, the response changed depending on the concentration of Reelin used (Fig 4A-B); the maximum effect was observed at a concentration of 30 nM similar to that previously used in mammalian hippocampal and cortical neurons [53, 54]. Several parameters highlighting the complexity of the neurite arborization were studied. For instance, Reelin treatment increased the maximum length that neurites can reach compared with mock-treated neurons (Fig 4C). Besides, Reelin exposure also increased the distance (from the soma) in which the maximum number of neurites is observed (critical value) (Fig 4D).

**Fig 4.**
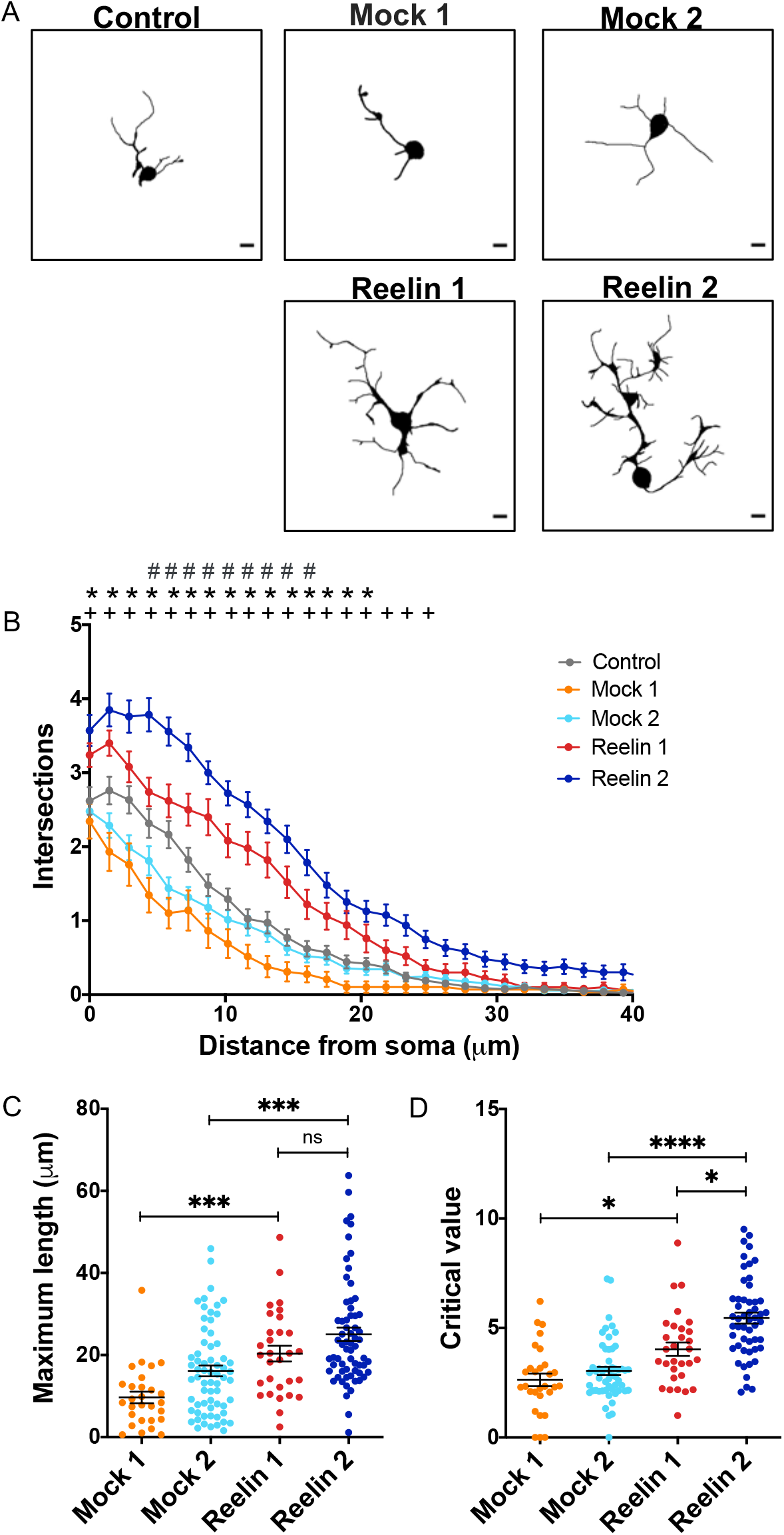
Mushroom body neurons respond to Reelin. **A** Representative images of MB neurons in primary cultures of the *c309,eGFP* strain, after Mock or Reelin treatment. 13 nM Reelin and 30 nM Reelin treatment correspond, respectively, to 20% and 50% dilution of Reelin conditioned medium in the cell culture. Mock 1 and Mock 2 treatments are the experimental controls of 13 nM and 30 nM Reelin, respectively. MB neurons are identified by GFP expression. Black bars in the lower right corner represent 5 μm. **B** Sholl profile of MB neurons after different treatments. Data are expressed as mean ± SEM; Two-way ANOVA, p<0.0001 for each factor. “*” indicates a significant difference (p<0.05) between the 13 nM Reelin and the corresponding mock 1 treatment, at a given distance from the soma; “+”, significant difference (p<0.05) between 30 nM Reelin and Mock 2 treatment, at the same distance from the soma. “#”, (p<0.05) between effects at 13 and 30 nM Reelin concentrations at the same distance from the soma. **C** Maximum length, defined as the greater distance reached for a cell’s neurite with respect to its soma, recorded in MB neurons under the different treatments. Data expressed as mean ± SEM; one-way ANOVA, followed by Kruskal Wallis post-hoc test, p<0.0001. *** indicates p<0.0005. **D** Critical value, the number of neurites found in the radius with most neurites, in MB neurons under each treatment. Data is mean ± SEM; one-way ANOVA and Kruskal Wallis post-test, p<0.0001. *, ****, are p<0.05, p<0.0001. “ns” not significant. Data from n=20 coverslips in four independent experiments, 15-20 cells in each “n”.

Next, we determined the role of LpRs in the morphological effects induced by Reelin in primary culture neurons. First, we verified the expression of LpRs in brain *Drosophila* cultured neurons (S6 Fig). Then, LpR1 or LpR2 deficient flies were used to prepare neuronal cultures that were exposed to 30 nM of Reelin. Our results showed that the LpRs-deficient MB neurons did not respond to Reelin, while responses to this signaling molecule were evident in cultured MB neurons from control animals (Fig 5). Interestingly, under basal conditions, LpR1 mutant neurons already showed impaired neurite development (Fig 5A). Other defects were also evidenced when studying the maximum length of neurites (Fig 5B) and the critical value (Fig 5C). Similarly, neuronal cultures derived from LpR2 mutant animals did not respond to Reelin, although a basal alteration in neurite development was not observed (Fig 5D-F).

**Fig 5.**
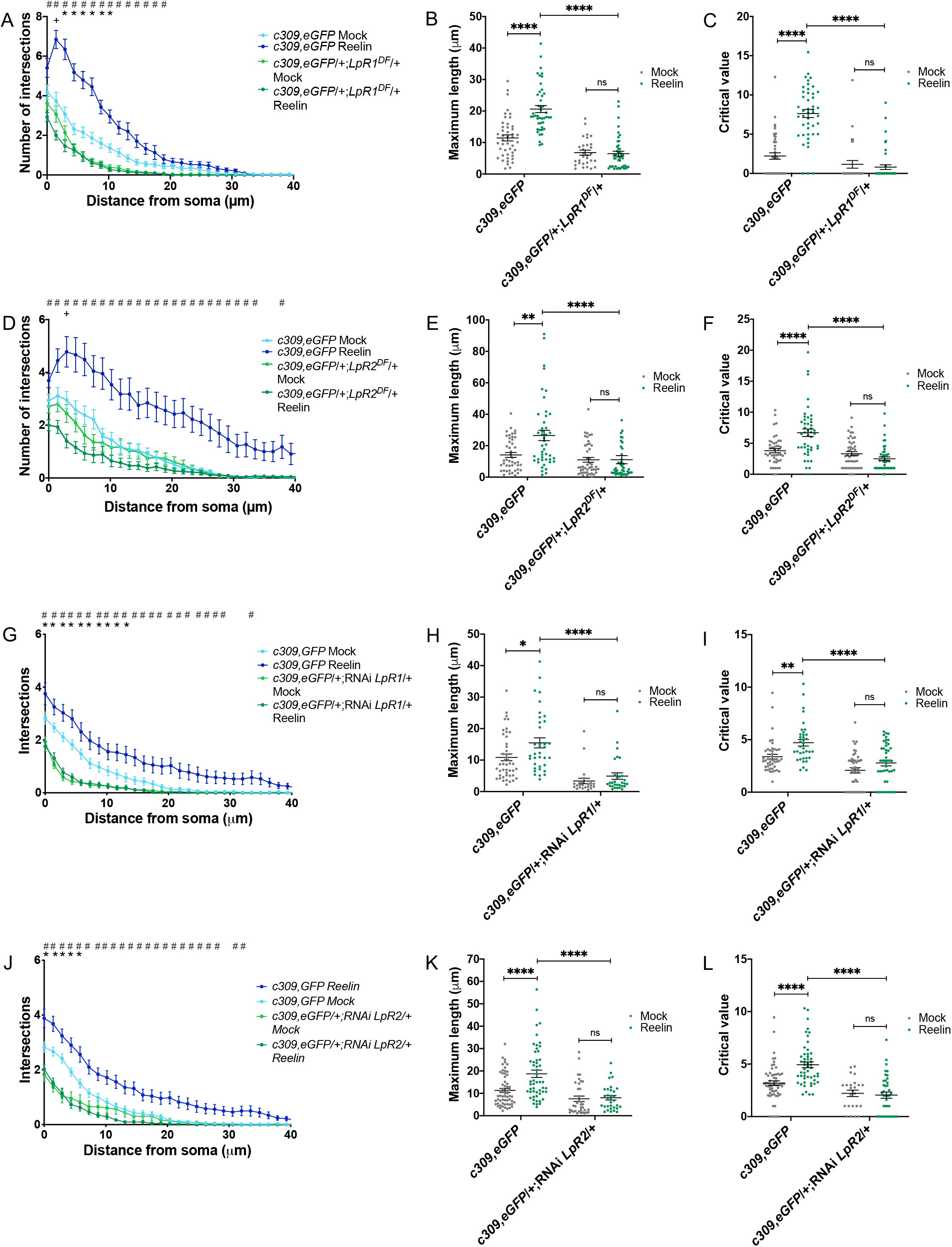
Reelin effects on mushroom body neurons depend on LpR1 and LpR2. **A-F** Reelin does not affect neurite complexity in mutants for LpRs. **A** and **D** Sholl profile of cultured MB neurons from LpR1 (*c309,eGFP*/+;*LpR1^DF^*/+) or LpR2 (*c309,eGFP*/+;*LpR2^DF^*/+) mutant flies and their controls (*c309,eGFP*), after Mock or Reelin treatment (30 nM). Two-way ANOVA shows that treatment and genotype factors and their interaction play a role in results obtained for each data set (p<0.0001 for all). “#” shows significant differences (p<0.05) between the genetic control and mutants after Reelin treatment. “*” indicates significant differences between mutant neurons and genetic control after Mock treatment. “+” means a significant difference (p<0.05) between mock and Reelin treatment in the mutant genotype. **B** and **E** Maximum length. **C** and **F** Critical value in the genotypes identified, after mock or Reelin treatment. In **B**, **C**, **E** and **F** Two-way ANOVA shows that treatment and genotype factors, and also the interaction between factors, play a role in results (p<0.0001 for each analysis); ** and ****, indicates p<0.01 and p<0.0001 between conditions. Data in **A**-**F**, from n=24 coverslips from three independent experiments, 15 cells studied in each coverslip. **G**-**L** Reelin does not affect neurite complexity LpRs knockdown flies. **G** and **J** Sholl profile, **H** and **K** Maximum length and **I** and **L** Critical value measured in flies expressing an RNAi for LpR1 or LpR2 only in MB neurons (*c309,eGFP*/+;RNAi-*LpR1*/+ or *c309,eGFP*/+;RNAi-*LpR2*/+, respectively), as compared to controls (*c309,eGFP*), after Mock or Reelin treatment. In **G** and **J** Two-way ANOVA shows that treatment and genotype factors and their interaction play a role in results obtained for each data set (p<0.0001 each). “#” shows significant differences (p<0.05) between control and LpR1 or LpR2 knockdown after Reelin treatment. “*” indicates significant differences between knockdown and control, after Mock treatment. No significant differences were observed between mock and Reelin treatment in the knockdowns for LpRs. In **H**, Two-way ANOVA indicates that only genotype and treatment factors play a role in results (p<0.0001 and p=0.0170 respectively). In **K**, two-way ANOVA indicates that genotype and treatment factors and their interaction play a role in results (p<0.0001, p=0.0034, and p=0.0096 respectively). In **I**, two-way ANOVA indicates that only genotype and treatment factors play a role in results (p<0.0001 and p=0.0002 respectively). In **L**, two-way ANOVA indicates that genotype and treatment factors and their interaction play a role in results (p<0.0001, p=0.0073, and p=0.0012 respectively). *, ** and **** indicated p<0.05, p<0.01 and p<0.0001 respectively. Data in **G**-**I**, from n=12 coverslips from three independent experiments, 10-15 cells studied in each coverslip. ns, not significant. Data in **J**-**L**, from n=16 coverslips from four independent experiments, 10-15 cells studied in each coverslip. ns, not significant.

To confirm these results and dissect the contribution of LpRs specifically in MB neurons, we took advantage of flies expressing RNAi for LpR1 or LpR2 only in the MB by using the Gal4-UAS system. By performing the same type of experiments outlined above, we obtained similar results to those described in cultured neurons from mutant flies: the inability of MB neurons to respond to Reelin when expressing RNAi for LpRs (Fig 5G-L). Overall, these results show for the first time that vertebrate Reelin stimulates the growth and neurite complexity of *Drosophila* MB neurons in a way that depends on the expression of LpRs.

### LpRs long isoforms are required to mediate Reelin internalization

The interaction of Reelin with ApoER2 and VLDL-R requires the LA modules in the extracellular domain of the receptors [14, 17], which are conserved in LpRs [2, 6]. Since the main difference between the different isoforms for LpRs is the presence of the first LA module plus the non-conserved domain, we asked whether the effect observed in neuronal cultures exposed to vertebrate Reelin depends on specific isoforms of LpRs. In order to work on this question, we used the insect cellular model S2, which does not express LpRs [9].

To test a functional Reelin-LpR interaction, we carried out a ligand internalization assay in which S2 cells were transiently transfected with the following available tools: two long isoforms of LpRs tagged with an HA epitope (i.e., LpR1J-HA or LpR2E-HA) and one short isoform of LpR2: LpR2F-HA (Fig 6A, Fig S4). Reelin was internalized only in cells expressing the longer forms of the receptors, either LpR1J-HA or LpR2E-HA (Fig 6A). Meanwhile, S2 cells transfected with the LpR2F isoform (Fig 6A) or the empty plasmid (S7 Fig A) did not internalize Reelin.

**Fig 6.**
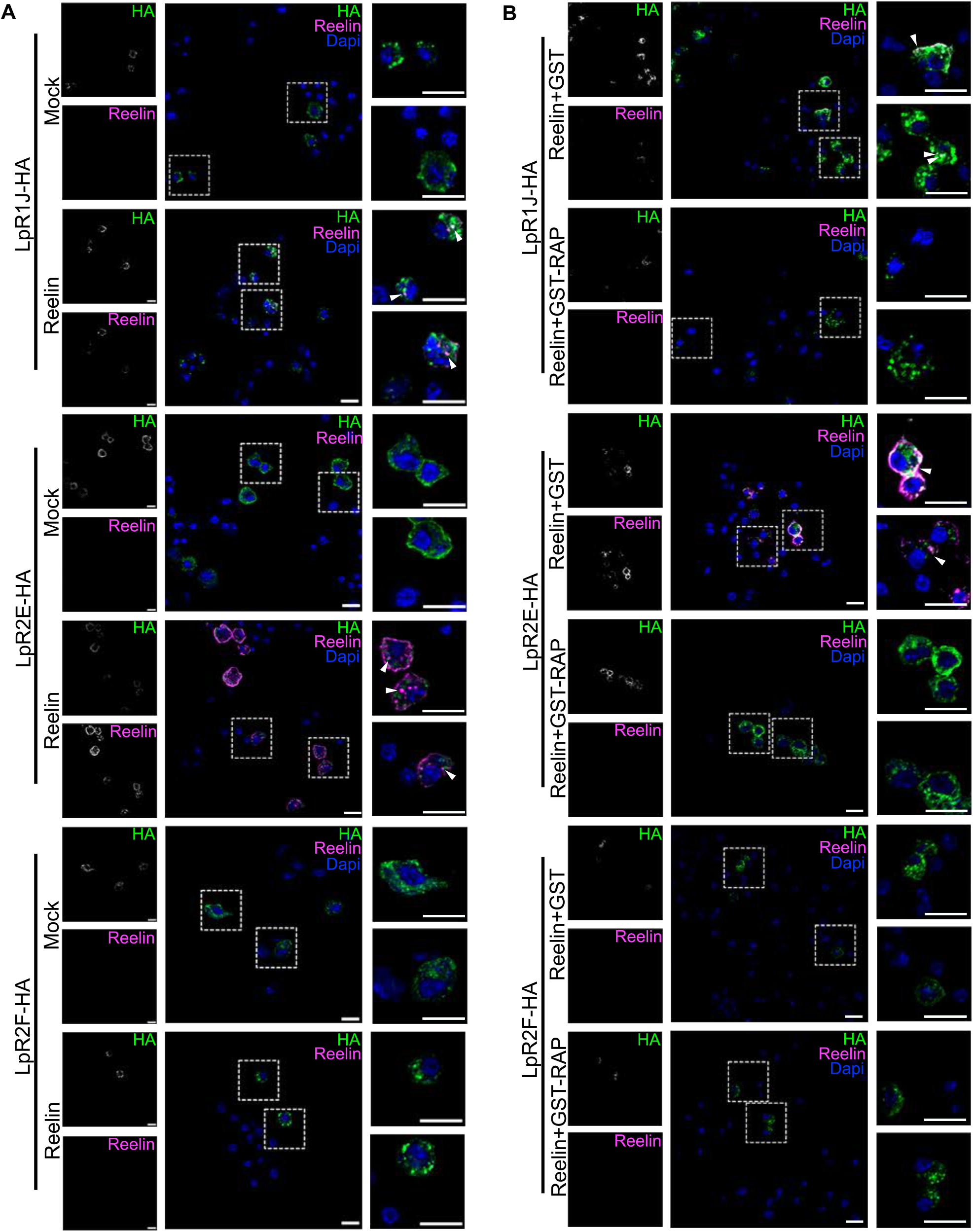
LpRs long isoforms internalize Reelin. **A** Representative images of S2 cells transfected either LpR1J-HA, LpR2E-HA or LpR2F-HA and treated with 30 nM Reelin or the equivalent volume of Mock media. Results of immunofluorescence against HA and Reelin. The panels in the right show magnifications of the area presented in the discontinuous white line square in the center image; data obtained from n=2 coverslips in one experiment. The lower right white bar in images represents 10μm. **B** Representative images of S2 cells transfected either LpR1J-HA, LpR2E-HA or LpR2F-HA and treated with 30 nM Reelin plus 500nM GST-RAP or 30nM Reelin plus 500nM GST. Results of immunofluorescence against HA and Reelin. The panels in the right show magnifications of the areas within the discontinuous white line square in the center image; data obtained from n=2 coverslips in one experiment. The lower right white bar represents 10μm.

The Receptor-Associated Protein (RAP) is a chaperone required for folding the LDL-R family members. RAP binds with high affinity to the ligand-binding domains of the receptors, preventing their premature interaction with ligands within the endoplasmic reticulum [55]. Several antecedents show that the extracellular addition of recombinant RAP prevents the binding of LDL-R family members to their respective ligands, including the binding of VLDL-R and ApoER2 with Reelin [28,55–57]. In addition, Van Hoof et al. [58] showed that the internalization of *Locusta migratoria* Lipophorin by its LpR could be completely inhibited by adding the recombinant form of the human RAP. Then, to evaluate if Reelin internalization depends on a classical interaction with lipoprotein receptors, the internalization experiment was performed in the presence of RAP (Fig 6B, S7 Fig B). Reelin entrance to the cells, mediated by LpR1J or LpR2E, was prevented by the presence of RAP in the media (Fig 6B).

Overall, these data suggest an interaction between mammalian Reelin and the two longer LpRs isoforms, which could underlie a novel, previously not described, signaling cascade.

### Dab is required for Reelin-induced responses in MB neurons

Reelin signaling requires the dimerization of ApoER2 or VLDL-R and the concomitant recruitment of the cytosolic adaptor protein Dab1 to the NPxY motif present in the cytoplasmic tail of these receptors [23,28,59]. In *Drosophila*, Disabled (Dab) is the homologous form of vertebrate Dab1 [26, 60]. To determine whether Dab contributes to the Reelin-induced response in MB neurons in culture, we used a mutant fly lacking one copy of *Dab.* Homozygous mutants for this gene die before reaching the third instar larvae [61]. The reduction of Dab expression in this mutant was corroborated by qPCR (S8 Fig A). We carried out the same type of *in vitro* experiments described when assessing the role of LpRs in primary neurons, but now in cultures prepared from *Dab* heterozygous mutants (Fig 7A-C). The Dab mutant neurons failed to respond to Reelin treatment: there was no change in neuronal arborization (Fig 7A), maximum length (Fig 7B), or the critical value (Fig 7C). As a complementary approach to corroborate these findings, Reelin was added to cultures prepared from brains expressing RNAi for Dab specifically in the MB, using c309-Gal4 (Fig 7D-F). The diminished expression of Dab was verified through qPCR (S8 Fig B). As with the Dab mutant cultures, Reelin treatment did not increase the complexity of neurite branching (Fig 7D), the maximum length (Fig 7E), or the critical value (Fig 7F) in MB neurons expressing RNAi for Dab. These results show that Dab plays a relevant role in the response of MB neurons to Reelin.

**Fig 7.**
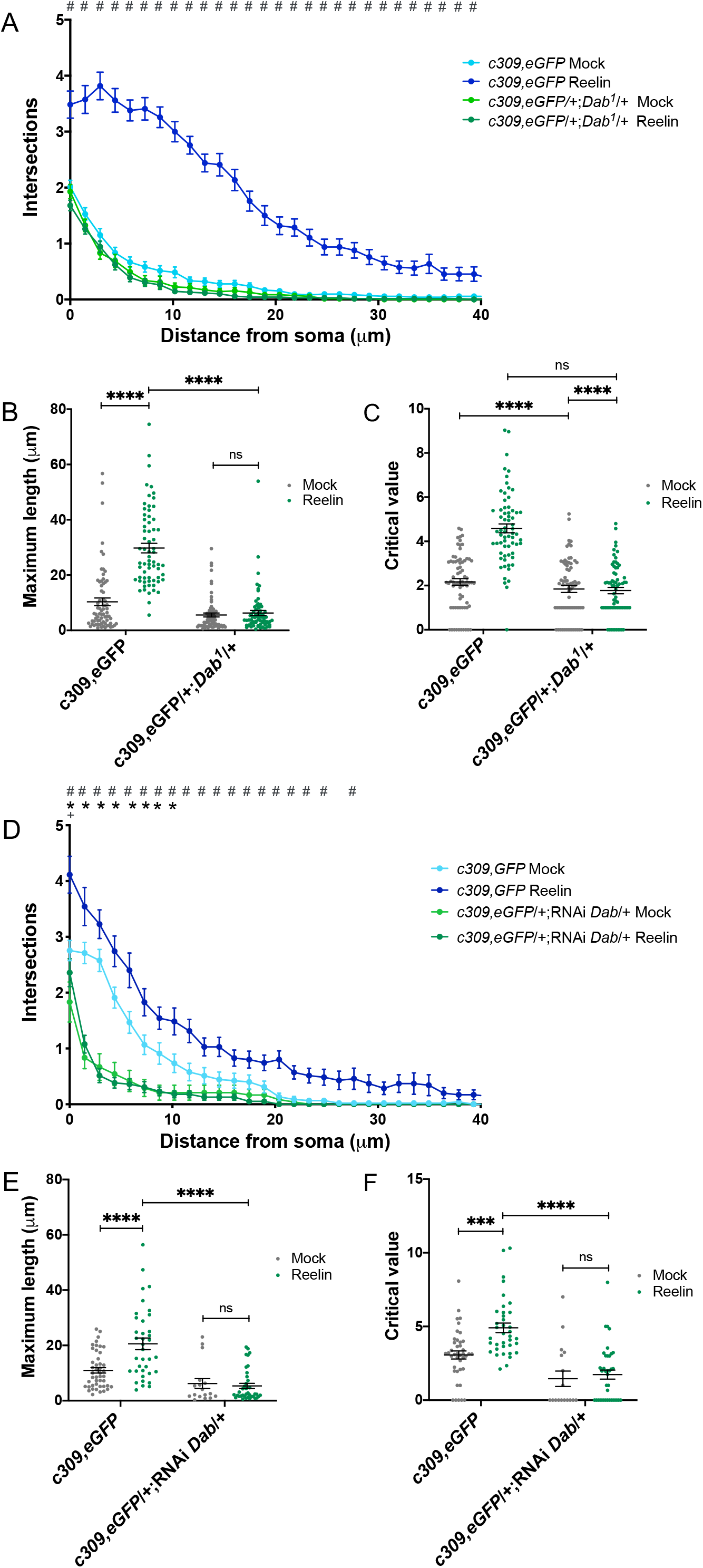
Reelin effects on mushroom body neurons depend on Dab. **A-C** Reelin does not increase neurite complexity in neurons from flies mutant for Dab. **A** Sholl profile of cultured MB neurons from Dab (*c309,eGFP*/+;*Dab^1^*/+) mutant flies and their controls (*c309,eGFP*), after Mock or Reelin treatment. Two-way ANOVA shows that treatment and genotype factors and their interaction play a role in results obtained for each data set (p<0.0001 for all). “#” shows significant differences (p<0.05) between the genetic control and mutant after they have been treated with Reelin. There were neither significant differences between mock and reelin treatment in the Dab mutant, nor between Mock treatment in the genetic control and Dab mutant. **B** Maximum length and **C** Critical value in mutant and control flies, after mock or Reelin treatment. In **B** and **C** two-way ANOVA shows that treatment and genotype factors and the interaction between factors play a role in the results (p<0.0001 for each analysis); ****, indicates p<0.0001 between conditions. Data in **A**-**C**, from n=12 coverslips obtained from three independent experiments, 15-20 cells studied in each coverslip. **D**-**F** Reelin does not affect neurite complexity in animals with knockdown for Dab. **D** Sholl profile, **E** Maximum length, and **F** Critical value measured in flies expressing an RNAi for Dab only in MB neurons (*c309,eGFP*/+;RNAi-*Dab*/+), as compared to controls (*c309,eGFP*), after Mock or Reelin treatment. Two-way ANOVA shows that treatment and genotype factors and their interaction play a role in results obtained for each data set (p<0.0001 for all). “#” shows significant differences (p<0.05) between control and Dab knockdown after Reelin treatment. “*” indicates significant differences between knockdown and control, after Mock treatment. “+” means a significant difference (p<0.05) between mock and reelin treatment on the fly that lacks a copy of Dab. In **E**, Two-way ANOVA indicates that genotype and treatment factors, and their interaction play a role in results (p<0.0001, p=0.0066, and p=0.0013 respectively). In **F** two-way ANOVA indicates genotype and treatment factors, and their interaction plays a role in results (p<0.0001, p=0.0031 and p=0.0285 respectively. In (E and F), *** and ****, p<0.005 and p<0.0001. “ns”, not significant. Data in **D**-**F**, from n=12 coverslips from three independent experiments, 4-15 cells studied in each coverslip.

### Dab is required for MB development and function

To determine whether mutant flies for Dab exhibit alterations in olfactory memory, we carried out the training protocol as outlined for *LpR* mutants. The performance index of flies lacking one copy of Dab was significantly lower than that of control animals (Fig 8A). These flies exhibited a normal response to the electric shocks and the odorants used (S9 Fig A-C). Additionally, we evaluated sleep patterns in animals deficient in Dab expression (Fig 8B and C). Mutant flies exhibited higher sleep time throughout the day, regardless of the phase (Fig 8B and C). Therefore, Dab would contribute to sleep homeostasis.

**Fig 8.**
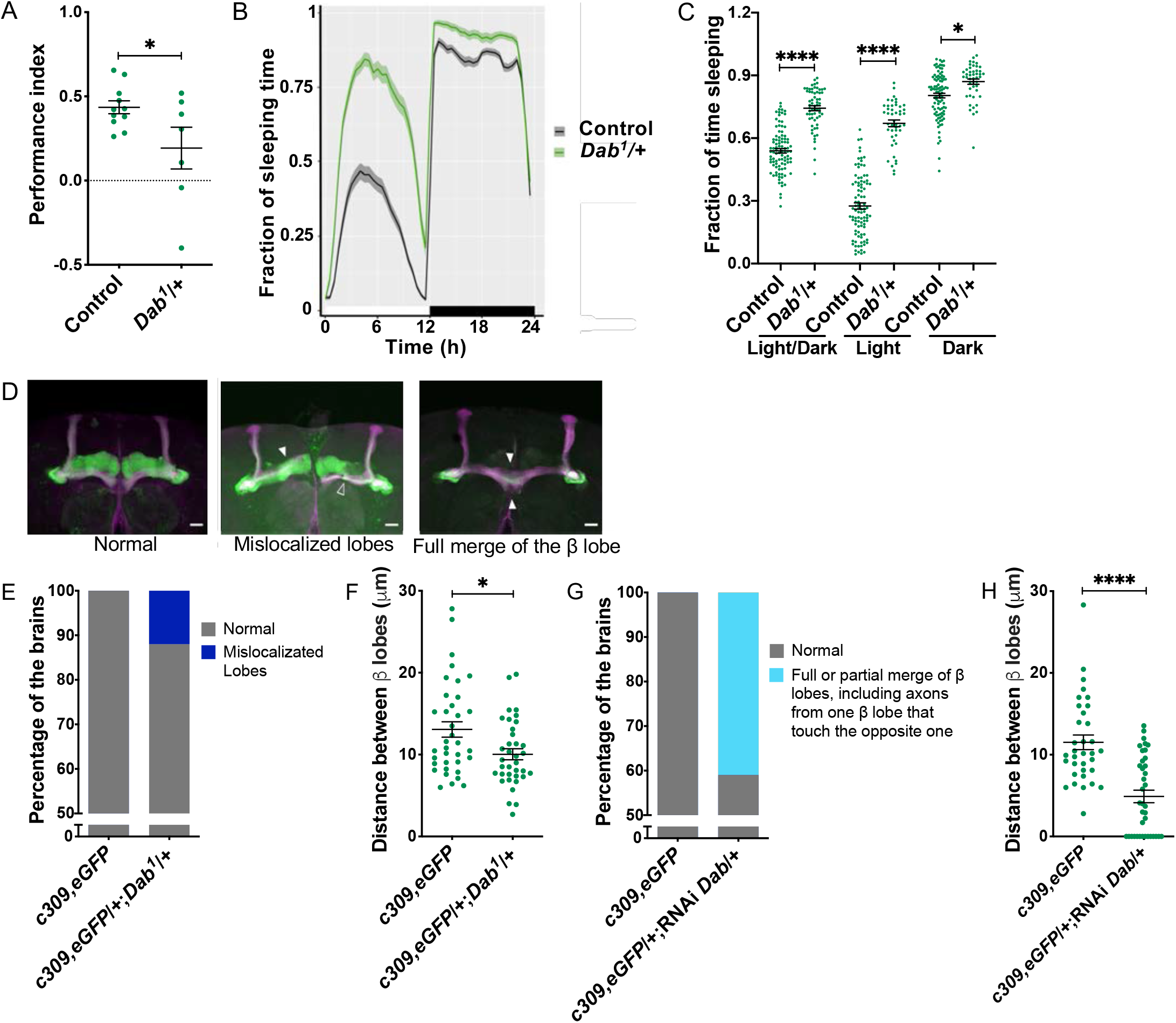
Dab is required for olfactory memory and establishing MB structure. **A** Memory performance is reduced in *Dab^1^* mutants as compared to control animals. Data is mean ± SEM; n= 7 independent experiments, t-test, p=0.0407 (* significant difference). **B** Sleep profile in *Dab* mutant flies. The white bar above the x-axis represents the hours of the day when flies were exposed to light; the black bar represents hours when flies were exposed to darkness. **C** Fraction of sleeping time in *Dab* mutants. Data in **B** and **C**, from n=2 independent experiments, 15-25 flies studied per genotype in each n. Data is mean ± SEM. Two-way ANOVA shows that light and genotype factors, and also the interaction between factors, contribute to results (p<0.0001 for each analysis), Tukey post-test; *, and ****, indicates p<0.05, and p<0.0001 between conditions. **D** Representative images of the phenotypes observed in MB of flies lacking one copy of Dab (*c309,eGFP*/+;*Dab^1^*/+) or express an RNAi against Dab in MB neurons (*c309,eGFP*/+; RNAi-*Dab*/+). Green, GFP; Magenta, Fas II immunostaining. Images are from several stacks. Empty arrowheads show normal MB structure; full arrowheads indicate an abnormality. White bar: 20 μm. **E** Distribution of phenotypes observed in control (n=4 independent experiments, 36 brains) and mutant (n=4 experiments, 34 brains) animals; Fisher test, p=3.44*10^-4^. **F** Distance between β lobes in brains from Dab mutants as compared to controls; t-test, shows significant differences (* p=0.0107); data in *c309,eGFP* from n=4 independent experiments, 36 brains in total and *c309,eGFP*/+;*Dab^1^*/+ from n=4 experiments, 40 brains. **G** Relative percentage for phenotypes observed when RNAi against Dab transcript is expressed in MB neurons. Fisher test, p=5.029*10^-15^ in *c309,eGFP* (n=3 experiments, 26 brains) and in *c309,eGFP*/+;*Dab^1^*/+ (n=3 experiments, 34 brains) flies. **H** Distance between β lobes in flies expressing RNAi for Dab in MB neurons as compared to control animals; t-test, shows significant differences (**** p<0.0001); data from *c309,eGFP* (n=4 experiments, 35 brains) and *c309,eGFP*/+;*Dab^1^*/+ (n=3, 38 brains) animals.

We also studied the contribution of Dab to MB organization by using two different genetic tools: mutants generated by a deletion in the Dab gene or animals expressing, only in MB neurons, an RNAi for transcripts of this gene (Fig 8D-H, S4 Table and S5 Table). The phenotypes observed in the Dab deficient animals included mislocalized lobes in 11 .7% of brains (Fig 8E). Besides, Dab mutants exhibit a lower distance between β lobes (Fig 8F). On the other hand, 41.2% of the brains from flies expressing RNAi against Dab exhibit MB with β lobes fused or axons crossing from one β lobe to the opposite (Fig 8G and H). Therefore, compared to controls, MB from Dab deficient flies exhibit their β lobes closer.

These results show that the presence of Dab is necessary for MB development and function in a similar way to LpRs.

## Discussion

Here we have demonstrated that LpR1, LpR2, and the adaptor protein Dab have a relevant and previously unnoticed role in the correct anatomical organization of adult MB and consequently in its associated functions, including olfactory memory performance and sleep patterns. Besides, the exposure of cultured *Drosophila* MB neurons to mammalian Reelin, a protein not present in flies, resulted in augmented neurite growth and complexity, an effect that depends on these receptors and Dab. Further, two of the LpRs long isoforms, LpR1J and LpR2E, are required for the internalization of Reelin in S2 cells.

### LpRs are required for adult MB structure and operation

The expression of LpR1 or LpR2 or the intracellular adapter Dab in MB was crucial for proper performance in memory assays. Further experiments are required to evaluate whether LpRs expressed in specific MB subpopulations differentially contribute to olfactory memory. The MB, fly brain region linked to the generation and storage of olfactory memories [37, 62], is proposed to play a homologous function to the mammalian hippocampus [63–66]. The MB and the hippocampus categorize stimuli according to the type of signal received and the context in which the signals are received, fundamental aspects for learning and memory processes [64]. Moreover, numerous ortholog genes are required in the hippocampus and MB for similar functions [66].

On the other hand, LpRs also participate in normal sleep homeostasis, another function associated with MB [33, 34]. Dab mutants also exhibited a higher sleep time throughout the day. Whether LpRs and Dab are required in brain regions beyond MB for sleep homeostasis is an issue we did not explore in this work. However, Yin et al. [12] showed that the reshuffle of LNvs dendrites under light stimuli was prevented when LpRs expression was eliminated [12]. Future studies will advance on the contribution of LpRs and Dab in sleep regulation.

The defects in MB structure observed in mutants for LpR1, LpR2 and Dab indicate the importance of these proteins in the appropriate organization of this fly brain region along with the development, determining its function in adult animals. Although we did not observe a high expression of LpR1 in the adult MB, our data show that it is strongly expressed in the β’ lobe in the third instar larvae, a stage where the MB cells are developing [48]. Our expression studies demonstrating diffuse expression of LpR1 and LpR2 in third instar larvae are coincident with previous reports [11, 12]. Interestingly, our results using the LpR1-Gal4 tool in adulthood and pupae stage showed that the receptor’s highest expression is not in the MB itself but in a group of cells surrounding the β’/β lobe of this structure. The complementary use of the LpR1 antibody supports the notion that these cells could be R neurons from the ellipsoid body [43–45].

Axonal patterning defects could underlie some phenotypes observed in animals deficient for LpRs [67]. For example, the fusion of the β−lobes is an indication of excessive and uncontrolled growth of axons crossing towards the contralateral hemisphere. It could be speculated that axons bearing LpRs and Dab should sense a stop signal close to the midline. When some intrinsic or extrinsic components are not present, the β−axons cross the midline. In this regard, it is worth highlighting a parallel with the role of ApoER2, VLDL-R, Dab1, and Reelin in the process of cell migration and axonal growth and targeting [68]. Reelin functions as an attractive cue for neuronal migration in the hippocampus and the cortex [19,49,69], as well as for axonal targeting in the visual system [70, 71] and the entorhinal-hippocampus circuit [72, 73]. Besides, Reelin behaves as a stop signal in the vicinity of cells secreting it (i.e., Cajal-Retzius in the developmental cortex and hippocampus). Therefore, in the absence of the receptors, Dab1 or Reelin, neurons invade the marginal zone [23, 60]. In the case of Reelin, the stop signal relies on the activation of Rho-GTPases and LIMK [22, 74]. In *Drosophila*, the growth of MB axons of the α/β lobes is also controlled by attractive and repulsive cues, and in some cases, as for the TGF-β 1 receptor, the stop signal includes LIMK activation [75].

Other phenotypes observed upon LpRs knockdown are stalling of MB lobes. Previous studies show that overexpression of LIMK and some manipulation of Rho GTPases activity lead to MB axon stalling [76, 77], which is consistent with the role of reelin in the control of these cytoskeleton regulators. Finally, defects in guidance in which the α and β lobes follow the same trajectory have been observed in mutants of the Eph-Ephrin [78] and Wnt5/Drl1/2 [79] signaling pathways, indicating LpR could be part of a complex signaling network regulating MB axon guidance.

Consistent with the structural and functional studies in the *Drosophila* brain, the experiments carried out in primary cultured neurons support the importance of LpRs in the structure of MB cells. Our data show impairment in basal neurite development in cultured neurons from mutants for these genes. The primary neuronal culture is a cell system generated from pupal brains, and therefore, phenotypes observed argue in favor of the idea that LpRs play a role in MB development.

Although the correlation between alterations in MB structure and behavioral phenotypes has been previously reported for several mutants of *Drosophila* genes [40, 80], to our understanding, this is the first time that this is shown for mutants in LpRs or the adaptor Dab. Overall, our results suggest that LpR and Dab regulate several aspects of MB morphogenesis and function, and future studies should address its interaction with other pathways involved in these processes.

### Hints on a novel function for the LpRs

Regarding the function of LpRs in *Drosophila*, both receptors have been primarily associated with lipid uptake [5,6,9,10]. Thus, disturbing LpR1 or LpR2 expression in early developmental stages reduces neutral lipid content in oocytes and imaginal discs. This phenotype is rescued by expressing long isoforms of LpRs, LpR1H and LpR2E [6]. ApoD and ApoE are endocytosed in mammalian cells by ApoER2 and VLDL-R [1, 81]. Recent work from Yin et al. [11] shows that a short isoform of LpR1 (LpR1G) interacts with the lipocalin Glaz. Interestingly, a report from the Bellen group shows that the transport of lipids between neurons and glia is mediated by Glaz and neural lazarillo proteins (Nlaz), the orthologs for ApoD in flies [82]. The mechanism associated with lipid uptake by the glia is unknown. It is then possible to hypothesize that some of the alterations we described in MB in LpRs-deficient animals could be due to deficits in the transport of lipids between neurons and glia. Interestingly, Matsuo et al. [9] showed that the lack of LpR1 and LpR2 does not significantly affect triglycerides levels in the CNS of third instar larvae, while a lower content was evidenced only for some phospholipids. This phenotype was also described by Yin et al. [11]. Since the highest expression for LpRs is described at the larval stage, the work of Matsuo et al. has several implications. First of all, it indicates that there are probably other complementary mechanisms responsible for the uptake of neutral lipids, so that deficiency on LpRs is not associated with a dramatic decrease in larval lipid content. Also, it supports the idea that LpRs could play other roles in addition to lipid uptake. In this regard, it was shown that LpR1 and LpR2 are required for proper LNv dendrite development and that these receptors are necessary for LNv structural plasticity in response to light stimuli in third instar larvae [12]. The mechanisms by which LpRs function would associate with increased synaptic activity were not discussed [12].

Strikingly, our data show that cultured MB neurons respond to Reelin treatment in a way depending on the expression of LpRs and Dab. The Reelin-induced increase in neurite arborization is similar to what has been previously described in cultured hippocampal neurons [50–53]. Likewise, the absence of a Reelin-induced response in cultured fly neurons deficient in LpRs is similar to the situation observed in cultured vertebrate neurons when the function of ApoER2 and VLDL-R is inhibited [52, 54].

Our results also show that the MB phenotypes observed in LpR1 mutants were rescued after expression in MB of the long isoform of LpR1, LpR1J, suggesting that the MB structure and anatomy of its constituent neurons depend on this LpR1 long isoform. Moreover, we found that this isoform binds and internalizes mammalian Reelin in *Drosophila* cells. Although re-expression in MB neurons of LpR2E, a long isoform of LpR2, was not effective at reverting the mutant structural phenotype observed in adult MB, our data showed that this isoform was also involved in Reelin internalization. Since we only evaluated some of the LpR isoforms, we cannot discard that other LpRs would mediate Reelin interaction and internalization or that other long isoforms participate in MB development. Nevertheless, our results suggest that LpRs short isoforms would have a different role than the two long isoforms.

The prevention of Reelin internalization by RAP indicates that Reelin would bind LpR1J and LpR2E isoforms through its cysteine-rich repeats in the LA module, like the rest of the LDL-R family members [6,55,83]. Reelin signaling also requires the interaction of Dab1 through its PTB domain with the NPxY motif of ApoER2 and VLDL-R [26,28,84]. As mentioned, the same motif is found in LpR1 and LpR2 [6], while the PTB domain is conserved in Dab from *Drosophila melanogaster* [26]. Furthermore, once Dab1 binds to ApoER2 or VLDL-R, it is phosphorylated in two groups of tyrosine residues [59], and it is already known that *Drosophila* Dab tyrosines can be phosphorylated [85]. The phosphorylation of Dab1 is crucial for the activation of downstream signaling molecules, including PI3K and Crk/CrkL, which results in the modification of cytoskeleton dynamics and membrane trafficking [1,20,22,50,51,86–89]. Thus, it would be interesting to evaluate, in the future, whether Reelin binding to LpRs triggers a signaling pathway that depends on the phosphorylation of Dab.

As previously mentioned, Reelin is not present in flies. However, a study suggests that protein domains in vertebrate Reelin appeared in evolution with the filum Arthropoda, to which *Drosophila* belongs [90], supporting that a Reelin-like protein could exist in at least some invertebrates. We are currently analyzing the *Drosophila melanogaster* genome to unveil whether flies have a protein with a homolog function to vertebrate Reelin.

## Conclusions

We have established new physiological and developmental roles for LpR1, LpR2, and Dab in *Drosophila* MB by using heterozygous mutants and RNAi targeted expression. Overall, all these results are consistent with the idea that we have uncovered a novel signaling pathway relevant to MB formation and function. Besides, our data provide support for the existence of a Reelin-like protein in *Drosophila*.

## Material and methods

### Drosophila stocks

Flies strains and F1 progeny when needed were raised under 12hrs:12hrs light:dark cycle at 25°C. The flies were fed with standard fly food. The flies employed were generated using flies from Bloomington Drosophila Stock Center (BDSC): y^1^w*; Mi{PT-GFSTF.1}^LpR2MI04745GFSTF.1^ (RRID:BDSC_60219), LpR1-Gal4: *w^1118^; PBac{w^+mC^=IT.GAL4}LpR1^0104-G4^/TM6B,Tb^1^* (RRID:BDSC_62639), *LpR1^DF^*: *w*;P{w^+mC^=XP-U}LpR1^Df^/TM6B,Tb^1^* (from RRID:BDSC_32653), RNAi LpR1: *w*; P{y^+t7.7^ v^+t1.8^=TRiP.JF02551}attP2 LpR1* (isogenized to *w^1118^* genetic background from RRID:BDSC_27249), *LpR2^DF^*: *w^1118^;rho^ve-1^ PBac{w^+mC^=RB3.WH3}LpR2^Df^/TM6B,Tb^1^* (from RRID:BDSC_44233), RNAi LpR2: *w*; P{y^+t7.7^ v^+t1.8^=TRiP.JF01627}attP2 LpR2* (isogenized to *w^1118^* genetic background from RRID:BDSC_31150), *Dab^1^*: *w^1118^;Dab^1^/TM6B, Tb^1^* (from RRID:BDSC_32653) and RNAi Dab: *w*;P{y^+t7.7^ v^+t1.8^=TRiP.HMS02482}attP2 Dab* (isogenized to *w^1118^* background from RRID:BDSC_42646). In addition, *Elav*-Gal4, *c309*-Gal4,UAS-*eGFP* (*c309,eGFP*), UAS*CD8::GFP*;;*OK107*-Gal4, *c309*-Gal4; UAS-*CD4::tdTomato, UAS-LpR1D-HA (gift from Dr. J. Culi), UAS-LpR1J-HA (gift from Dr. J. Culi), UAS-LpR2E-HA (gift from Dr. J. Culi), UAS-LpR2FHA (gift from Dr. J. Culi)* were used.

### Measurement of mRNA levels

To demonstrate the reduction in the expression of mRNA levels in strains of interest, 50 adult male heads (0-3 days) were homogenized in TRIzol (Invitrogen). Chloroform (Merck) was added, mixed, and then centrifuged at 12,000 rpm (4°C) for 15 min. The aqueous phase was collected, and then mixed with isopropanol (Merck), followed by incubation, and posterior centrifugation at 12,000 rpm (4°C) for 20 minutes. The pellet was air-dried and then resuspended in nuclease-free water at 55°C for 15 min. RNA integrity was evaluated by running an aliquot of this preparation in an agarose gel, and RNA concentration was determined in an Epoch microplate reader (Biotek). The cDNA synthesis was carried out from 3 μg of RNA using the kit RevertAid First Strand cDNA Synthesis (Invitrogen).

The primers were directed to two exons that are present in all isoforms of the genes: LpR1: 5’–TGCACGAATGGAGCCTGCAT–3’ and 5’–GTGATGCGATCCTTGCACTGGT–3’, annealing at 60°C. LpR2: 5’–CTATGTCGTATACCGACGCTGC–3’ and 5’–CTGCGGCGTAAATGTGTGG–3’ annealing at 57°C. Dab: 5’– GTCTTAGCACCACGAATGGAA–3’ and 5’– GGCGTTATCCGTTCCATCT–3’ annealing at 55°C. GAPDH was used as a reference gene: 5’ – CGTTCATGCCACCACCGCTA–3’ and 5’– CCACGTCCATCACGCCACAA–3’ annealing at 60°C. The reaction used the 5x HOTFIREPol® EvaGreen® qPCR Mix Plus Kit (Solis BioDyne), and LightCycler® capillaries (20 μL), in the LightCycler® (Roche) apparatus. Primer efficiency was evaluated after serial dilution and was calculated as 10^-1/slope^. The relative expression was calculated as [primer efficiency for GAPDHCT average among replica]/ [primer efficiency for the gene of interest CT average among replica].

Each MB is composed of about ∼2500 neurons [36] out of the about 100,000 found in fly brains. To test the RNAi effectiveness by qPCR, the RNAi of interest was pan-neurally driven by Elav-Gal4 (Fig S1C and D, Fig S8B). This is a gal4 driver for the whole nervous system, and it is not expressed in other cell types in the fly head. Then the expression measured corresponds to an estimation of the mRNA levels.

### Aversive olfactory conditioning

Groups of 40-50 flies were collected and kept at 25°C and 70% relative humidity under 12 hr:12hr light/dark cycles in the behavioral room. Experiments were conducted using a T-maze apparatus connected to a constant airflow (3 liters/min) to provide flies with either fresh air or a given odorant. The process consists of airflow passing through bubbling mineral oil or a solution of the odorant dissolved in mineral oil. The experiments were conducted under dim red light to prevent confounding visual effects on olfactory memory formation [91].

On the day of the experiment, flies were transferred into a training tube lined with an electrified grid. After 90 s acclimatization to the tube, flies were exposed to an odorant (the conditioned stimulus, CS^+^), either 3-octanol (OCT) or 4-methyl cyclohexanol (MCH), paired to twelve 70 V DC electric shocks (the unconditioned stimulus, US) over 60 s (Fig 3). This procedure was followed by a 45 s rest period with fresh air. Flies were then exposed to the reciprocal odorant (CS^-^), which is not paired to electric shock. Memory was evaluated one h post-training to test Mid-term memory (MTM). For this, flies are exposed to both odorants, each in one tube; the number of flies choosing each tube was quantified. A performance index (PI) was calculated as:

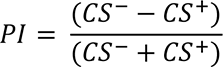

The CS^+^ odorant was reversed in alternate groups of flies, between OCT and MCH, to account for innate bias toward one odorant. Thus, n = 1 corresponds to the average of the PI of two consecutive alternated groups. Sensory-motor controls were performed to assess whether avoidance of electric shocks or odorants were similar between different strains. For olfactory acuity, the odorant and air were pumped through opposite arms of the T-maze; flies were allowed to choose between the two tubes for 2 min. For electric shock avoidance, flies were allowed to decide between two training tubes, one of them connected to the stimulator providing electric shock. For olfactory and shock processing, the fly avoidance over the stimuli was quantified, and responses were reported as the percentage of flies that avoided the stimuli over the total number of flies (S2 Fig).

### Locomotor and sleep analysis

Locomotor activity was measured using the Drosophila Activity Monitoring System (DAM, TriKinetics Inc, USA), using 7-9 days old male flies. The individuals were collected, separated by phenotype and transferred into DAM tubes containing standard fly food and placed into an incubator (25°C and 75% humidity) connected to the monitoring system (TriKinetics, Waltham, MA, United States) under 12 h:12 h light-dark cycles for 6–7 days. After six days of baseline recordings, the sleep analysis was performed in R using the Rethomics framework [92]. Sleep is defined as a continuous period of inactivity lasting 5 min or more [29, 30], and activity counts to actograms profiles (S2 Fig E and S9 Fig D) were calculated as the number of beam crossings each 30min.

### Fly brain immunohistochemistry

Brains from i) c309,eGFP/+;LpR1^DF^/+; ii) c309,eGFP/+;RNAi-LpR1/+; iii) c309,eGFP/+;LpR2^DF^ /+; and iv) c309,eGFP/+;RNAi-LpR2/+ 0-3 days old adult male flies were fixed for 20 minutes with 4% PFA, 4% Sucrose in PBS1X. The tissue was blocked (solution: 3% Normal Goat Serum (NGS), 0.03% Sodium Azide in Phosphate-buffered saline with 0.3 % Triton-X100 (PBS-T) for 1 hr, and then incubated with 1D4-FasII antibody (RRID: AB_528235) 1:20 at 4°C, overnight (ON). After three hours of incubation with secondary antibody Alexa Fluor 568 goat anti mouse (RRID: AB_144696) 1:500, at room temperature (RT), brains were mounted with ProLong Gold with DAPI (Invitrogen).

### Cell culture immunocytochemistry

Primary cultured neurons were fixed with 4% Paraformaldehyde (PFA) and 4% sucrose in PBS1X for 20 minutes. Then, cell cultures were incubated for 15 minutes with 0.15M Glycine and washed three times with PBS-T. The cells were blocked (solution: 3% NGS, 0.03% Azide in PBS-T) for 1 hr. followed by overnight (ON) incubation at 4° with the primary antibodies: Guinea Pig anti LpR1 (Gifted by J. Culi) 1:100 or GFP antibody (RRID: AB 94936) 1:500. Then, the coverslips were washed three times with PBST and incubated for 2 hrs. at RT with secondary antibodies. Secondary antibodies used were goat anti-guinea pig 648 (RRID: AB_2340476) 1:350 or Alexa Goat anti-mouse 488 (RRID: AB_2633275). Later, the cells were washed three times with PBS-T. The coverslips were mounted in Fluoromount-G with DAPI.

### Preparation of primary cultures of pupal brain neurons

As previously reported [93, 94]. In brief, fly brains were dissected out of the animals at pupal stage P9, in dissecting solution (DS), consisting of 6.85 mM NaCl, 0.27 mM KCl, 0.009 mM Na2HPO4, 0.001 mM KH2PO4, 0.2772 mM HEPES, pH 7.4. After enzymatic treatment of the tissue (9.6 U/ml of papain, Worthington, LS 03126) 30 minutes, at room temperature, mechanical disaggregation was carried out in culture plates containing Laminin-Concanavalin A (Sigma) coated coverslips in the presence of DMEM/F12 culture medium (Gibco, 12400-016) supplemented with 100μg/ml Apo-Transferrin, 30 nM Selenium, 50 μg/ml Insulin, 2.1 mM 20-Hydroxyecdysone, 20 ng/ml Progesterone, 100 μM Putrescine (all supplements from Sigma-Aldrich), and 1% Antibiotic/Antimycotic (Gibco 15240062). The following day, the cells received conditioned media obtained from astrocytes cultured in Neurobasal medium supplemented with B27 (CNBM/27).

### Reelin production

Reelin was obtained as conditioned media from a cell line (HEK293T) that stably expresses mammalian Reelin (D’Arcangelo et al., 1997). Cells were cultured in DMEM high glucose (Gibco) supplemented with Fetal Bovine Serum (10% FBS, Biological Industries), 1% Penicillin/Streptomycin (Gibco) and G418 (0.5 mg/ml). In order to collect Reelin, the culture media is serum-free. The supernatant containing Reelin was collected for five days, and the medium was concentrated using Amicon ultra-15 100kDa filters (Millipore). Reelin concentration in the medium was estimated semiquantitatively: BSA dilution curves and an aliquot of Reelin-containing medium were run through an SDS-PAGE. The gel was stained with Coomassie blue to visualize reelin bands. After that, the bands corresponding to BSA and Reelin were digitalized to analyze intensities in Fiji [95].

### Reelin treatments for Sholl Analysis

Primary neuronal cultures were prepared on 12 mm coverslips and maintained in 24well plates. Five days in vitro (DIV) cultures were treated with Reelin at 13 or 30 nM for 48 hrs. In parallel, neurons were treated with culture media collected from Hek293 cells transfected with the empty vector (Mock medium). Afterward, cultures are fixed and mounted, as mentioned above.

### Internalization assay

S2 cells were transfected with plasmids pAC5.1-LpR1J-HA, pAC5.1-LpR2E-HA, pAC5.1-LpR2F-HA, or the empty vector using the calcium phosphate method. After 48 h of expression, the cells were treated with 30 nM Reelin or the equivalent volume of Mock media for 30 min at 27°C. Then, the cells were washed one time with Schneider medium, twice with the same medium but at pH=3.2, and two more times with Schneider medium. The cells were fixed, and immunocytochemistry was performed as mentioned above. The primary antibodies used were HA antibody (RRID: AB_1549585) 1:500 and E4 Reelin antibody (RRID: AB_1157891) 3:17. The secondary antibodies were Alexa Fluor 488 Chicken anti Rabbit (RRID: AB_2535859) 1:500 and Alexa Fluor 568 goat anti-mouse (RRID: AB_144696) 1:500.

## Analyses

MB morphology was evaluated in both hemispheres in Fiji. It also measured the distance between the MB lobes in Fiji [95].

Statistical analyses were performed using GraphPad Prism, except for the Fisher test, which was performed using R and “fisher. test” from package “stats” version 4.0.1.

## List of abbreviations

LpRs: Lipophorin Receptors
LpR1: Lipophorin Receptor 1
LpR2: Lipophorin Receptor 2
MB: Mushroom Bodies
LDL-R: Low-Density Lipoprotein Receptor
ApoE: Apolipoprotein E
ApoB: Apolipoprotein B
LNv: ventral Lateral Neurons
CNS: central nervous system
Dab1: Disabled 1
CS+: conditioned stimulus
US: unconditioned stimulus
RAP: Receptor-Associated Protein
Dab: Disabled.

## Declarations

### Ethics approval

All procedures were approved by Comité de Ética y Seguridad, Pontificia Universidad Católica de Chile (ID Protocol: 190806006)

### Consent for publication

Not applicable.

### Availability of data and materials

The datasets used and/or analyzed during the current study are available from the corresponding author on reasonable request.

### Contributions

M-P.M, JMC and F.R-C conceived and designed the study. The formal analysis where developed by F.R-C., N.F-U, and SH. F.R-C., V.T-V., N.F-U. SH, M-C.G-R. and CBR conducted the experiments. JMC, M-P.M. and CO. provided the resources for the study. F.R-C. wrote the original manuscript. M-P.M., JMC, F.R-C. and CO. reviewed & edited the manuscript.

## Acknowledgments

We are very grateful to Dr. Joaquim Culi (Universidad Autónoma de Madrid, Spain) for providing us the antibodies to detect LpR1 and the flies and the plasmids to express the different isoforms of LpRs. We are indebted to Dr. Joachim Herz (University of Texas Southwestern, USA), who kindly provided the HEK293 Reelin-producing cells. We also thank Gloria Loyola for her technical support and Dr. Estela Andrés for accessing the RT-qPCR apparatus. Many thanks to Dr. James Hodge (University of Bristol, UK) for giving us access to the olfactory learning and memory apparatus and Dr. Alfredo Ghezzi (Universidad de Puerto Rico, Rio Piedras) for allowing us to use the circadian setup. We acknowledge the help provided by the Advanced Microscopy Facility UC.

## Supporting information

**S1 Fig. Verification of reduction in LpR1 or LpR2 expression in mutant and knockdown animals. Related to Figs 1, 2, and 5. A-B** Reduced expression of LpR1 or LpR2 in flies heterozygous for deletion in *LpR1* or *LpR2* genes (*c309,eGFP*/+; *LpR1^DF^*/+ and *c309,eGFP*/+;*LpR2^DF^*/+, respectively) evaluated through qPCR. Data from n=5 independent experiments for each genotype; * and ** means significant differences after paired t-test, p=0.0128 and p=0.0018, respectively. **C**-**D** Reduced LpR1 and LpR2 expression in flies expressing RNAi for each transcript (*Elav(x);;*UAS-RNAi *LpR1* and *Elav(x)*;;UAS-RNAi *LpR2*) in the whole nervous system by using the Elav driver. Data from n=4 and 5 independent experiments, respectively; * significant differences after paired t-test (p=0.0391 and p=0.0439)

**S2 Fig. Sensory controls in LpR1 and LpR2 mutant flies. Related to Fig 1. A** Response to electric shocks in flies that lack a copy of LpR1 or LpR2 compared to control. Data expressed as mean ± SEM; n=4 independent experiments, one-way ANOVA P=0.9687, no differences between groups. **B** Innate response to MCH. Data expressed as mean ± SEM of n=4 independent experiments, one-way ANOVA (P=0.0201) followed by Kruskal Wallis test. Flies mutant for LpR1 exhibit a higher sensitivity to MCH than controls; * P<0.05, “ns” no significant differences. **C** Innate response to Octanol (Oct). Data expressed as mean ± SEM from n=4 independent experiments. One-way ANOVA shows no differences between groups (p=0.1082). **D** Comparison in performance index in Cs^w-^ and *w^1118^* flies; t-test p=0.1743, no differences between genotypes. Data expressed as mean ± SEM from n= 11 or 3 independent experiments, respectively. **E** Locomotor activity profile throughout the day in flies mutant for LpR1 or LpR2. Data from n=2, two independent experiments, 15-25 flies studied per genotype in each n. Data expressed as mean ± SEM. Two-way ANOVA, Tukey post-test, shows that the hour of the day, genotype factors, and the interaction between factors contribute to results (p<0.0001 for each analysis). “+” indicates a significant difference (p<0.05) between Control fly and mutant for LpR1, at a given time of the day; “#”, a significant difference (p<0.05) between Control and LpR2 mutant flies at the same hour of the day.

**S1 Table. Phenotypes of flies lacking one copy of LpR1 or LpR2 and after the re-expression of specific LpRs isoforms. Related to Fig 2.**

**S2 Table. Phenotypes of flies that express RNAi against LpR1 or LpR2 in MB neurons under the control of the c309-Gal4 driver. Related to Fig 2**.

**S3 Fig. Knocking down LpR1 or LpR2 in MB neurons using the OK107 driver results in altered MB structure. Related to Fig 2. A** Percentage of brains that exhibit different phenotypes after expression of RNAi for LpR1 or LpR2 in MB, driven by OK107-Gal4. Data from CD8::GFP/+;;OK107 flies (n=3 experiments, 25 brains altogether); CD8::GFP/+;RNAiLpR1/+;OK107/+ animals (n=3 experiments, 24 brains in total); CD8::GFP/+;RNAi LpR2/+;OK107/+ flies (n=3 experiments, 29 brains); Fisher test, p=2.2*10-16. **B** Distance between β lobes after knocking down LpR1 or LpR2 in the MB by using the OK107 driver. Data expressed as mean ± SEM from CD8::GFP/+;;OK107 (n=3 experiments, 26 brains); CD8::GFP/+;RNAi-LpR1/+;OK107/+ (n=3 experiments, 26 brains) and CD8::GFP/+;RNAiLpR2/+;OK107/+ (n=3 experiments, 29 brains) flies. One-way ANOVA (p<0.0001) followed by Kruskal Wallis. ***, **** means p<0.0005 and p<0.0001 respectively. See also Table Supplementary 3. “ns” means not statically significant.

**S3 Table. Phenotypes of flies expressing RNAi against LpR1 or LpR2 in MB neurons under the control of the OK107 driver. Related to Fig S3.**

**S4 Fig. Schematic representation of LpRs isoforms studied.** Organization of the domains in the LpRS isoforms studied with its correspondence exons for LpR1 (**A**) and LpR2 (**B**). At the top of each panel, the green box represents the exons labels 1-15. The dark green mark indicates the UTR regions for each cDNA.

**S5 Fig. LpR1 expression in *W^1118^* adulthood flies. Related to Fig 3.** Representative image of immunofluorescence against LpR1 and FasII in *w^1118^* flies with DAPI staining. The discontinuous line in the merge indicates the magnified region in the inset. Scale white bars represent 50 μm.

**S6 Fig. LpRs are expressed in primary culture neurons from pupal brain. Related to Fig 5. A** Representative results of immunofluorescence experiment to detect LpR1 in primary cultures of a *Drosophila* brain where MB neurons express the membrane protein CD8-ChRFP under the control of the c309-Gal4 driver (c309-Gal4>UAS-CD8.ChRFP). Arrowheads show MB-positive cells identified by CD8.ChRFP expression. **B** Representative images of immunofluorescence staining for GFP in primary cultures of a *Drosophila* brain expressing LpR2-GFP^MI04745^ where MB neurons are identified by CD8.ChRFP expression. Arrowheads indicate MB-positive cells as identified in. The white bar represents 5 μm.

**S7 Fig. Control of the Reelin internalization. Related to Fig 6. A** Representative images of immunocytochemistry of anti-HA and Reelin in S2 cells transiently transfected with the empty vector pAC5.1 which were treated with 30nM Reelin or the equivalent volume of Mock media under the same internalization protocol as in Fig 6A. **B** Representative images of immunocytochemistry of anti-HA and Reelin in S2 cells transiently transfected with the empty vector pAC5.1 which were treated with 30nM Reelin plus 500nM of GST-RAP or 500nM GST under the same internalization protocol used in Fig 6B.

**S8 Fig. Verification of reduction in Dab expression in mutant and knockdown animals. Related to Figs 7 and 8. A** Reduced expression of Dab in mutant *c309,eGFP*/+;*Dab^1^*/+ flies lacking one copy of the gene evaluated through qPCR. Data expressed as mean ± SEM; paired t-test, * p=0.0105. Data from n=5 independent experiments, 50 fly heads each “n”. **B** Expressing RNAi for Dab pan-neurally (*Elav(x)*/+;;UASRNAi *Dab*/+) results in decreased expression of transcript for the protein. Data from n=4 independent experiments for each genotype, 50 fly heads each “n”. * means statistical differences after paired t-test, p=0.0141.

**S9 Fig. Sensory controls in Dab mutant flies. Related to Fig 8. A** Shock avoidance in Dab mutants (*Dab^1^*/+). Data expressed as mean ± SEM of n=4 independent experiments; T-test p=0.1331, no significant differences. **B**-**C** Innate response to MCH and Oct, respectively, in mutant *Dab^1^*/+ flies compared to controls. Data expressed as mean ± SEM of n=4 independent experiments; T-test p=0.4860 and p=0.3429, no differences between groups. **D** Locomotor activity profile throughout the day in flies mutant for *Dab*. Data from n=2, two independent experiments, 15-25 flies studied per genotype in each n. Data expressed as mean ± SEM. Two-way ANOVA, Tukey post-test, showing that the hour of the day, genotype factors, and interaction between factors, play a role in results (p<0.0001 for each analysis). “#” indicates a significant difference (p<0.05) between Control and Dab mutant files at the same hour of the day. **E** Number of eggs from Dab mutant animals. T-test P=0.0065. ** mean P<0.01 n=3, from three independent experiments carried out with 30 females from *c309,eGFP*/+ and *c309,eGFP*/+;*Dab^1^*/+ strains. **F** Number of eggs from LpR1 and LpR2 mutants. One Way ANOVA, Tukey post-test P=0.0162 *means P<0.005. n=3, from three independent experiments carried out with 30 females from *c309,eGFP*/+;*LpR1^DF^*/+ and *c309,eGFP*/+;*LpR2^DF^*/+, and *c309,eGFP* control animals.

**S4 Table. Phenotypes of mutant flies lacking a copy of Dab. Related to Fig 8.**

**S5 Table. Phenotypes of flies that express an RNAi against Dab in MB neurons directed by *c309-Gal4*. Related to Fig 8.**

## References

1. Lane-Donovan C, Herz J. ApoE, ApoE Receptors, and the Synapse in Alzheimer’s Disease. Trends Endocrinol Metab. 2017;28: 273–284. https://doi.org/10.1016/j.tem.2016.12.001

2. Rodenburg K, Smolenaars M, Vanhoof D, Vanderhorst D. Sequence analysis of the non-recurring C-terminal domains shows that insect lipoprotein receptors constitute a distinct group of LDL receptor family members. Insect Biochem Mol Biol. 2006;36: 250– 263. https://doi.org/10.1016/j.ibmb.2006.01.003

3. Go G, Mani A. Low-density Lipoprotein receptor (LDLR) Family orchestrates cholesterol Homeostasis. Yale J Biol Med. 2012;85: 19–28.

4. Zanoni P, Velagapudi S, Yalcinkaya M, Rohrer L, von Eckardstein A. Endocytosis of lipoproteins. Atherosclerosis. 2018;275: 273–295. https://doi.org/10.1016/j.atherosclerosis.2018.06.881

5. Rodenburg K, Van der Horst D. Lipoprotein-mediated lipid transport in insects: Analogy to the mammalian lipid carrier system and novel concepts for the functioning of LDL receptor family members. Biochim Biophys Acta BBA - Mol Cell Biol Lipids. 2005; S1388198105001538. https://doi.org/10.1016/j.bbalip.2005.07.002

6. Parra-Peralbo E, Culi J. Drosophila Lipophorin Receptors Mediate the Uptake of Neutral Lipids in Oocytes and Imaginal Disc Cells by an Endocytosis-Independent Mechanism. Ashrafi K, editor. PLoS Genet. 2011;7: e1001297. https://doi.org/10.1371/journal.pgen.1001297

7. Soukup SF, Culi J, Gubb D. Uptake of the Necrotic Serpin in Drosophila melanogaster via the Lipophorin Receptor-1. Rulifson E, editor. PLoS Genet. 2009;5: e1000532. https://doi.org/10.1371/journal.pgen.1000532

8. Huang R, Song T, Su H, Lai Z, Qin W, Tian Y, et al. High-fat diet enhances starvation-induced hyperactivity via sensitizing hunger-sensing neurons in Drosophila. eLife. 2020;9: e53103. https://doi.org/10.7554/eLife.53103

9. Matsuo N, Nagao K, Suito T, Juni N, Kato U, Hara Y, et al. Different mechanisms for selective transport of fatty acids using a single class of lipoprotein in Drosophila. J Lipid Res. 2019;60: 1199–1211. https://doi.org/10.1194/jlr.M090779

10. Rodríguez-Vázquez M, Vaquero D, Parra-Peralbo E, Mejía-Morales JE, Culi J. Drosophila Lipophorin Receptors Recruit the Lipoprotein LTP to the Plasma Membrane to Mediate Lipid Uptake. Thummel CS, editor. PLOS Genet. 2015;11: e1005356. https://doi.org/10.1371/journal.pgen.1005356

11. Yin J, Spillman E, Cheng ES, Short J, Chen Y, Lei J, et al. Brain-specific lipoprotein receptors interact with astrocyte derived apolipoprotein and mediate neuron-glia lipid shuttling. Nat Commun. 2021;12: 2408. https://doi.org/10.1038/s41467-021-22751-7

12. Yin J, Gibbs M, Long C, Rosenthal J, Kim HS, Kim A, et al. Transcriptional Regulation of Lipophorin Receptors Supports Neuronal Adaptation to Chronic Elevations of Activity. Cell Rep. 2018;25: 1181–1192.e4. https://doi.org/10.1016/j.celrep.2018.10.016

13. Sepp KJ, Hong P, Lizarraga SB, Liu JS, Mejia LA, Walsh CA, et al. Identification of Neural Outgrowth Genes using Genome-Wide RNAi. Takahashi JS, editor. PLoS Genet. 2008;4: e1000111. https://doi.org/10.1371/journal.pgen.1000111

14. Andersen OM, Benhayon D, Curran T, Willnow TE. Differential Binding of Ligands to the Apolipoprotein E Receptor 2. Biochemistry. 2003;42: 9355–9364. https://doi.org/10.1021/bi034475p

15. Benhayon D, Magdaleno S, Curran T. Binding of purified Reelin to ApoER2 and VLDLR mediates tyrosine phosphorylation of Disabled-1. Mol Brain Res. 2003;112: 33–45. https://doi.org/10.1016/S0169-328X(03)00032-9

16. D’Arcangelo G, Homayouni R, Keshvara L, Rice DS, Sheldon M, Curran T. Reelin Is a Ligand for Lipoprotein Receptors. Neuron. 1999;24: 471–479. https://doi.org/10.1016/S0896-6273(00)80860-0

17. Yasui N, Nogi T, Takagi J. Structural Basis for Specific Recognition of Reelin by Its Receptors. Structure. 2010;18: 320–331. https://doi.org/10.1016/j.str.2010.01.010

18. Beffert U, Weeber EJ, Durudas A, Qiu S, Masiulis I, Sweatt JD, et al. Modulation of Synaptic Plasticity and Memory by Reelin Involves Differential Splicing of the Lipoprotein Receptor Apoer2. Neuron. 2005;47: 567–579. https://doi.org/10.1016/j.neuron.2005.07.007

19. D’Arcangelo, Graham G. Miao, Shu-Cheng Chen, Holly D. Scares, James I. Morgan, Tom Curran. A protein related to extracellular matrix proteins deleted in the mouse mutant reeler. Nature. 1995;374: 719–723. https://doi.org/10.1038/374719a0

20. Knuesel I. Reelin-mediated signaling in neuropsychiatric and neurodegenerative diseases. Prog Neurobiol. 2010;91: 257–274. https://doi.org/10.1016/j.pneurobio.2010.04.002

21. Ranaivoson FM, von Daake S, Comoletti D. Structural Insights into Reelin Function: Present and Future. Front Cell Neurosci. 2016;10. https://doi.org/10.3389/fncel.2016.00137

22. Santana J, Marzolo M-P. The functions of Reelin in membrane trafficking and cytoskeletal dynamics: implications for neuronal migration, polarization and differentiation. Biochem J. 2017;474: 3137–3165. https://doi.org/10.1042/BCJ20160628

23. Trommsdorff M, Gotthardt M, Hiesberger T, Shelton J, Stockinger W, Nimpf J, et al. Reeler/Disabled-like Disruption of Neuronal Migration in Knockout Mice Lacking the VLDL Receptor and ApoE Receptor 2. Cell. 1999;97: 689–701. https://doi.org/10.1016/S0092-8674(00)80782-5

24. Weeber EJ, Beffert U, Jones C, Christian JM, Förster E, Sweatt JD, et al. Reelin and ApoE Receptors Cooperate to Enhance Hippocampal Synaptic Plasticity and Learning. J Biol Chem. 2002;277: 39944–39952. https://doi.org/10.1074/jbc.M205147200

25. Howell BW. Mouse disabled (mDab1): a Src binding protein implicated in neuronal development. EMBO J. 1997;16: 121–132. https://doi.org/10.1093/emboj/16.1.121

26. Kawasaki F, Iyer J, Posey LL, Sun CE, Mammen SE, Yan H, et al. The DISABLED protein functions in CLATHRIN-mediated synaptic vesicle endocytosis and exoendocytic coupling at the active zone. Proc Natl Acad Sci. 2011;108: E222–E229. https://doi.org/10.1073/pnas.1102231108

27. Yun M, Keshvara L, Park C-G, Dickerson JB, Zheng J, Rock CO, et al. Crystal Structures of the Dab Homology Domains of Mouse Disabled 1 and 2. J Biol Chem. 2003;278: 36572–36581. https://doi.org/10.1074/jbc.M304384200

28. Hiesberger T, Trommsdorff M, Howell BW, Goffinet A, Mumby MC, Cooper JA, et al. Direct Binding of Reelin to VLDL Receptor and ApoE Receptor 2 Induces Tyrosine Phosphorylation of Disabled-1 and Modulates Tau Phosphorylation. Neuron. 1999;24: 481–489. https://doi.org/10.1016/S0896-6273(00)80861-2

29. Hendricks JC, Finn SM, Panckeri KA, Chavkin J, Williams JA, Sehgal A, et al. Rest in Drosophila Is a Sleep-like State. Neuron. 2000;25: 129–138. https://doi.org/10.1016/S0896-6273(00)80877-6

30. Shaw PJ. Correlates of Sleep and Waking in Drosophila melanogaster. Science. 2000;287: 1834–1837. https://doi.org/10.1126/science.287.5459.1834

31. Haynes PR, Christmann BL, Griffith LC. A single pair of neurons links sleep to memory consolidation in Drosophila melanogaster. eLife. 2015;4: e03868. https://doi.org/10.7554/eLife.03868

32. Martin J-R, Ernst R, Heisenberg M. Mushroom Bodies Suppress Locomotor Activity in Drosophila melanogaster. Learn Mem. 1998;5: 13.

33. Pitman JL, McGill JJ, Keegan KP, Allada R. A dynamic role for the mushroom bodies in promoting sleep in Drosophila. Nature. 2006;441: 753–756. https://doi.org/10.1038/nature04739

34. Sitaraman D, Aso Y, Jin X, Chen N, Felix M, Rubin GM, et al. Propagation of Homeostatic Sleep Signals by Segregated Synaptic Microcircuits of the Drosophila Mushroom Body. Curr Biol. 2015;25: 2915–2927. https://doi.org/10.1016/j.cub.2015.09.017

35. Vogt K, Schnaitmann C, Dylla KV, Knapek S, Aso Y, Rubin GM, et al. Shared mushroom body circuits underlie visual and olfactory memories in Drosophila. eLife. 2014;3: e02395. https://doi.org/10.7554/eLife.02395

36. Aso Y, Grübel K, Busch S, Friedrich AB, Siwanowicz I, Tanimoto H. The Mushroom Body of Adult Drosophila Characterized by GAL4 Drivers. J Neurogenet. 2009;23: 156– 172. https://doi.org/10.1080/01677060802471718

37. Heisenberg M. Mushroom body memoir: from maps to models. Nat Rev Neurosci. 2003;4: 266–275. https://doi.org/10.1038/nrn1074

38. Crittenden JR, Skoulakis EMC, Han K-A, Kalderon D, Davis RL. Tripartite Mushroom Body Architecture Revealed by Antigenic Markers. Learn Mem. 1998;5: 38–51.

39. Huang C, Zheng X, Zhao H, Li M, Wang P, Xie Z, et al. A Permissive Role of Mushroom Body α/β Core Neurons in Long-Term Memory Consolidation in Drosophila. Curr Biol. 2012;22: 1981–1989. https://doi.org/10.1016/j.cub.2012.08.048

40. Pascual A. Localization of Long-Term Memory Within the Drosophila Mushroom Body. Science. 2001;294: 1115–1117. https://doi.org/10.1126/science.1064200

41. Zwarts L, Vanden Broeck L, Cappuyns E, Ayroles JF, Magwire MM, Vulsteke V, et al. The genetic basis of natural variation in mushroom body size in Drosophila melanogaster. Nat Commun. 2015;6: 10115. https://doi.org/10.1038/ncomms10115

42. Connolly JB, Roberts IJH, Armstrong JD, Kaiser K, Forte M, Tully T, et al. Associative Learning Disrupted by Impaired Gs Signaling in Drosophila Mushroom Bodies. Science. 1996;274: 2104–2107. https://doi.org/10.1126/science.274.5295.2104

43. Martín-Peña A, Acebes A, Rodríguez J-R, Chevalier V, Casas-Tinto S, Triphan T, et al. Cell types and coincident synapses in the ellipsoid body of *Drosophila*. Eur J Neurosci. 2014;39: 1586–1601. https://doi.org/10.1111/ejn.12537

44. Omoto JJ, Nguyen B-CM, Kandimalla P, Lovick JK, Donlea JM, Hartenstein V. Neuronal Constituents and Putative Interactions Within the Drosophila Ellipsoid Body Neuropil. Front Neural Circuits. 2018;12: 103. https://doi.org/10.3389/fncir.2018.00103

45. Zhang Z, Li X, Guo J, Li Y, Guo A. Two Clusters of GABAergic Ellipsoid Body Neurons Modulate Olfactory Labile Memory in Drosophila. J Neurosci. 2013;33: 5175–5181. https://doi.org/10.1523/JNEUROSCI.5365-12.2013

46. Nagarkar-Jaiswal S, Lee P-T, Campbell ME, Chen K, Anguiano-Zarate S, Cantu Gutierrez M, et al. A library of MiMICs allows tagging of genes and reversible, spatial and temporal knockdown of proteins in Drosophila. eLife. 2015;4: e05338. https://doi.org/10.7554/eLife.05338

47. Kunz T, Kraft KF, Technau GM, Urbach R. Origin of Drosophila mushroom body neuroblasts and generation of divergent embryonic lineages. Development. 2012;139: 2510–2522. https://doi.org/10.1242/dev.077883

48. Lee T, Lee A, Luo L. Clonal analysis of the mushroom bodies. Development. 1999;126: 4065–4076.

49. D’Arcangelo G. Reelin in the Years: Controlling Neuronal Migration and Maturation in the Mammalian Brain. Adv Neurosci. 2014;2014: 1–19. https://doi.org/10.1155/2014/597395

50. Jossin Y, Goffinet AM. Reelin Signals through Phosphatidylinositol 3-Kinase and Akt To Control Cortical Development and through mTor To Regulate Dendritic Growth. Mol Cell Biol. 2007;27: 7113–7124. https://doi.org/10.1128/MCB.00928-07

51. Matsuki T, Pramatarova A, Howell BW. Reduction of Crk and CrkL expression blocks reelin-induced dendritogenesis. J Cell Sci. 2008;121: 1869–1875. https://doi.org/10.1242/jcs.027334

52. Niu S, Renfro A, Quattrocchi CC, Sheldon M, D’Arcangelo G. Reelin Promotes Hippocampal Dendrite Development through the VLDLR/ApoER2-Dab1 Pathway. Neuron. 2004;41: 71–84. https://doi.org/10.1016/S0896-6273(03)00819-5

53. Sotelo P, Farfán P, Benitez ML, Bu G, Marzolo M-P. Sorting Nexin 17 Regulates ApoER2 Recycling and Reelin Signaling. Wanjin H, editor. PLoS ONE. 2014;9: e93672. https://doi.org/10.1371/journal.pone.0093672

54. Ampuero E, Jury N, Härtel S, Marzolo M-P, van Zundert B. Interfering of the Reelin/ApoER2/PSD95 Signaling Axis Reactivates Dendritogenesis of Mature Hippocampal Neurons: Reelin signaling maintains adult network stability. J Cell Physiol. 2017;232: 1187–1199. https://doi.org/10.1002/jcp.25605

55. Bu G, Marzolo MP. Role of RAP in the Biogenesis of Lipoprotein Receptors. 2000;10: 148–155. https://doi.org/10.1016/S1050-1738(00)00045-1

56. Battey FD, Gåfvels ME, FitzGerald DJ, Argraves WS, Chappell DA, Strauss JF, et al. The 39-kDa receptor-associated protein regulates ligand binding by the very low density lipoprotein receptor. J Biol Chem. 1994;269: 23268–23273. https://doi.org/10.1016/S0021-9258(17)31648-4

57. Stockinger W, Hengstschläger-Ottnad E, Novak S, Matus A, Hüttinger M, Bauer J, et al. The Low Density Lipoprotein Receptor Gene Family. J Biol Chem. 1998;273: 32213–32221. https://doi.org/10.1074/jbc.273.48.32213

58. Van Hoof D. Insect lipoprotein follows a transferrin-like recycling pathway that is mediated by the insect LDL receptor homologue. J Cell Sci. 2002;115: 4001–4012. https://doi.org/10.1242/jcs.00113

59. Keshvara L, Benhayon D, Magdaleno S, Curran T. Identification of Reelin-induced Sites of Tyrosyl Phosphorylation on Disabled 1. J Biol Chem. 2001;276: 16008–16014. https://doi.org/10.1074/jbc.M101422200

60. Howell BW, Hawkes R, Soriano P, Cooper JA. Neuronal position in the developing brain is regulated by mouse disabled-1. Nature. 1997;389: 733–737. https://doi.org/10.1038/39607

61. Song JK, Kannan R, Merdes G, Singh J, Mlodzik M, Giniger E. Disabled is a bona fide component of the Abl signaling network. Development. 2010;137: 3719–3727. https://doi.org/10.1242/dev.050948

62. McGuire SE. The Role of Drosophila Mushroom Body Signaling in Olfactory Memory. Science. 2001;293: 1330–1333. https://doi.org/10.1126/science.1062622

63. Hourcade B, Muenz TS, Sandoz JC, Rossler W, Devaud JM. Long-Term Memory Leads to Synaptic Reorganization in the Mushroom Bodies: A Memory Trace in the Insect Brain? J Neurosci. 2010;30: 6461–6465. https://doi.org/10.1523/JNEUROSCI.0841-10.2010

64. Maza FJ, Sztarker J, Shkedy A, Peszano VN, Locatelli FF, Delorenzi A. Context-dependent memory traces in the crab’s mushroom bodies: Functional support for a common origin of high-order memory centers. Proc Natl Acad Sci. 2016;113: E7957– E7965. https://doi.org/10.1073/pnas.1612418113

65. Owald D, Waddell S. Olfactory learning skews mushroom body output pathways to steer behavioral choice in Drosophila. Curr Opin Neurobiol. 2015;35: 178–184. https://doi.org/10.1016/j.conb.2015.10.002

66. Wolff GH, Strausfeld NJ. Genealogical correspondence of a forebrain centre implies an executive brain in the protostome–deuterostome bilaterian ancestor. Philos Trans R Soc B Biol Sci. 2016;371: 20150055. https://doi.org/10.1098/rstb.2015.0055

67. Stoeckli ET. Understanding axon guidance: are we nearly there yet? Development. 2018;145: dev151415. https://doi.org/10.1242/dev.151415

68. Sacks D, Baxter B, Campbell BCV, Carpenter JS, Cognard C, Dippel D, et al. Multisociety Consensus Quality Improvement Revised Consensus Statement for Endovascular Therapy of Acute Ischemic Stroke. Int J Stroke. 2018;13: 612–632. https://doi.org/10.1177/1747493018778713

69. Del Río JA, Heimrich B, Borrell V, Förster E, Drakew A, Alcántara S, et al. A role for Cajal–Retzius cells and reelin in the development of hippocampal connections. Nature. 1997;385: 70–74. https://doi.org/10.1038/385070a0

70. Di Donato V, De Santis F, Albadri S, Auer TO, Duroure K, Charpentier M, et al. An Attractive Reelin Gradient Establishes Synaptic Lamination in the Vertebrate Visual System. Neuron. 2018;97: 1049–1062.e6. https://doi.org/10.1016/j.neuron.2018.01.030

71. Rice DS, Nusinowitz S, Azimi AM, Martínez A, Soriano E, Curran T. The Reelin Pathway Modulates the Structure and Function of Retinal Synaptic Circuitry. Neuron. 2001;31: 929–941. https://doi.org/10.1016/S0896-6273(01)00436-6

72. Borrell V, Del Río JA, Alcántara S, Derer M, Martínez A, D’Arcangelo G, et al. Reelin Regulates the Development and Synaptogenesis of the Layer-Specific Entorhino-Hippocampal Connections. J Neurosci. 1999;19: 1345–1358. https://doi.org/10.1523/JNEUROSCI.19-04-01345.1999

73. Borrell V, Pujadas L, Simó S, Durà D, Solé M, Cooper JA, et al. Reelin and mDab1 regulate the development of hippocampal connections. Mol Cell Neurosci. 2007;36: 158–173. https://doi.org/10.1016/j.mcn.2007.06.006

74. Chai X, Förster E, Zhao S, Bock HH, Frotscher M. Reelin acts as a stop signal for radially migrating neurons by inducing phosphorylation of n-cofilin at the leading edge. Commun Integr Biol. 2009;2: 375–377. https://doi.org/10.4161/cib.2.4.8614

75. Ng J. TGFβ signals regulate axonal development through distinct Smad-independent mechanisms. Development. 2008;135: 4025–4035. https://doi.org/10.1242/dev.028209

76. Ng J, Luo L. Rho GTPases Regulate Axon Growth through Convergent and Divergent Signaling Pathways. Neuron. 2004;44: 779–793. https://doi.org/10.1016/j.neuron.2004.11.014

77. Ng J, Nardine T, Harms M, Tzu J, Goldstein A, Sun Y, et al. Rac GTPases control axon growth, guidance and branching. Nature. 2002;416: 442–447. https://doi.org/10.1038/416442a

78. Boyle M, Nighorn A, Thomas JB. Drosophila Eph receptor guides specific axon branches of mushroom body neurons. Development. 2006;133: 1845–1854. https://doi.org/10.1242/dev.02353

79. Reynaud E, Lahaye LL, Boulanger A, Petrova IM, Marquilly C, Flandre A, et al. Guidance of Drosophila Mushroom Body Axons Depends upon DRL-Wnt Receptor Cleavage in the Brain Dorsomedial Lineage Precursors. Cell Rep. 2015;11: 1293– 1304. https://doi.org/10.1016/j.celrep.2015.04.035

80. Heisenberg M, Borst A, Wagner S, Byers D. The Mushroom Body Mutants are Deficient in Olfactory Learning. J Neurogenet. 1985;2: 1–30. https://doi.org/10.3109/01677068509100140

81. Kosacka J, Gericke M, Nowicki M, Kacza J, Borlak J, Spanel-Borowski K. Apolipoproteins D and E3 exert neurotrophic and synaptogenic effects in dorsal root ganglion cell cultures. Neuroscience. 2009;162: 282–291. https://doi.org/10.1016/j.neuroscience.2009.04.073

82. Liu L, MacKenzie KR, Putluri N, Maletić-Savatić M, Bellen HJ. The Glia-Neuron Lactate Shuttle and Elevated ROS Promote Lipid Synthesis in Neurons and Lipid Droplet Accumulation in Glia via APOE/D. Cell Metab. 2017;26: 719–737.e6. https://doi.org/10.1016/j.cmet.2017.08.024

83. Tufail M, Takeda M. Insect vitellogenin/lipophorin receptors: Molecular structures, role in oogenesis, and regulatory mechanisms. J Insect Physiol. 2009;55: 88–104. https://doi.org/10.1016/j.jinsphys.2008.11.007

84. Howell BW, Lanier LM, Frank R, Gertler FB, Cooper JA. The Disabled 1 Phosphotyrosine-Binding Domain Binds to the Internalization Signals of Transmembrane Glycoproteins and to Phospholipids. Mol Cell Biol. 1999;19: 5179– 5188. https://doi.org/10.1128/MCB.19.7.5179

85. Gertler FB, Hill KK, Clark MJ, Hoffmann FM. Dosage-sensitive modifiers of Drosophila abl tyrosine kinase function: prospero, a regulator of axonal outgrowth, and disabled, a novel tyrosine kinase substrate. Genes Dev. 1993;7: 441–453. https://doi.org/10.1101/gad.7.3.441

86. Beffert U, Morfini G, Bock HH, Reyna H, Brady ST, Herz J. Reelin-mediated Signaling Locally Regulates Protein Kinase B/Akt and Glycogen Synthase Kinase 3β. J Biol Chem. 2002;277: 49958–49964. https://doi.org/10.1074/jbc.M209205200

87. Chen K. Interaction between Dab1 and CrkII is promoted by Reelin signaling. J Cell Sci. 2004;117: 4527–4536. https://doi.org/10.1242/jcs.01320

88. Howell BW, Herrick TM, Hildebrand JD, Zhang Y, Cooper JA. Dab1 tyrosine phosphorylation sites relay positional signals during mouse brain development. Curr Biol. 2000;10: 877–885. https://doi.org/10.1016/S0960-9822(00)00608-4

89. Leemhuis J, Bouche E, Frotscher M, Henle F, Hein L, Herz J, et al. Reelin Signals through Apolipoprotein E Receptor 2 and Cdc42 to Increase Growth Cone Motility and Filopodia Formation. J Neurosci. 2010;30: 14759–14772. https://doi.org/10.1523/JNEUROSCI.4036-10.2010

90. Manoharan M, Muhammad SA, Sowdhamini R. Sequence Analysis and Evolutionary Studies of Reelin Proteins. Bioinforma Biol Insights. 2015;9: BBI.S26530. https://doi.org/10.4137/BBI.S26530

91. Tully T, Quinn WG. Classical conditioning and retention in normal and mutant Drosophila melanogaster. J Comp Physiol [A]. 1985;157: 263–277. https://doi.org/10.1007/BF01350033

92. Geissmann Q, Garcia Rodriguez L, Beckwith EJ, Gilestro GF. Rethomics: An R framework to analyse high-throughput behavioural data. Flack JC, editor. PLOS ONE. 2019;14: e0209331. https://doi.org/10.1371/journal.pone.0209331

93. Leyton V, Goles N. I., Fuenzalida-Uribe N., Campusano J. M. V. Octopamine and Dopamine differentially modulate the nicotine-induced calcium response in Drosophila Mushroom Body Kenyon Cells. Neuroscience Letters. 2014;560: 16–20. http://dx.doi.org/10.1016/j.neulet.2013.12.006

94. Sicaeros B, Campusano JM, O’Dowd DK. Primary Neuronal Cultures from the Brains of Late Stage Drosophila Pupae. J Vis Exp. 2007; 200. https://doi.org/10.3791/200

95. Schindelin J, Arganda-Carreras I, Frise E, Kaynig V, Longair M, Pietzsch T, et al. Fiji: an open-source platform for biological-image analysis. Nat Methods. 2012;9: 676–682. https://doi.org/10.1038/nmeth.2019

