## Supplementary Information for "Lipophorin receptors regulate mushroom bodies development and participate in learning, memory and sleep in flies"

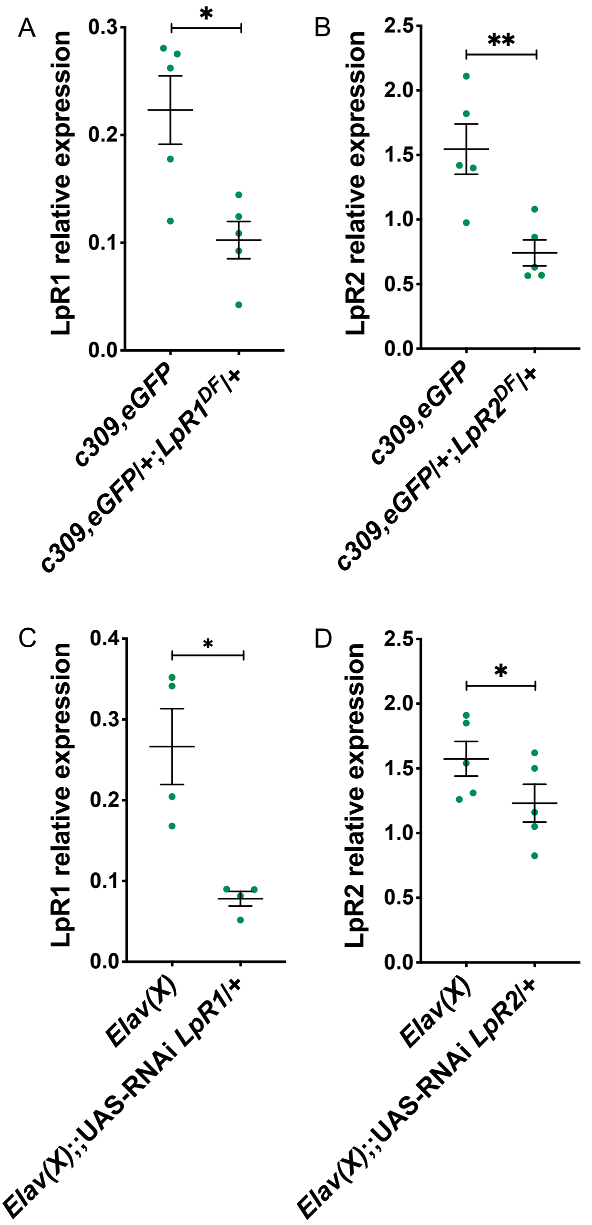


**S1 Fig. Verification of reduction in LpR1 or LpR2 expression in mutant and knockdown animals. Related to Figs 1, 2 and 5.**

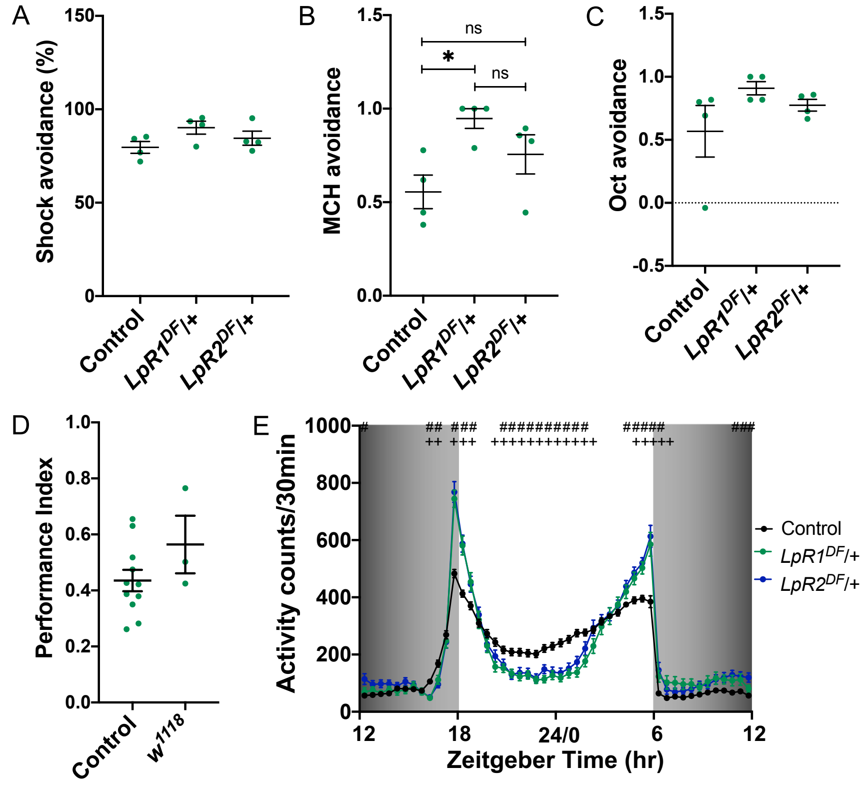


**S2 Fig. Sensory controls in LpR1 and LpR2 mutant flies. Related to Fig 1.**

**S1 Table. Phenotypes of flies lacking one copy of LpR1 or LpR2 and after the re-expression of specific LpRs isoforms. Related to Fig 2.**

| | ***c309,eGFP*** | ***c309,eGFP*/+; *LpR1$^{DF}$*/+** | ***c309,eGFP*/UAS-LpR1D-HA; *LpR1$^{DF}$*/+** | ***c309,eGFP*/UAS-LpR1J-HA; *LpR1$^{DF}$*/+** | ***c309,eGFP*/+; *LpR2$^{DF}$*/+** | ***c309,eGFP*/UAS-LpR2E-HA; *LpR2$^{DF}$*/+** | ***c309,eGFP*/UAS-LpR2F-HA; *LpR2$^{DF}$*/+** |
|---|---|---|---|---|---|---|---|
| Normal | 40 | 34 | 20 | 19 | 43 | 26 | 20 |
| Emergence of axons from a β lobe that touch the opposite β lobe | | | 1 | | 2 | 1 | 1 |
| merge of β lobes | | 1 | 1 | | | | |
| Mislocalizated α | | 2 | | | | | |
| Mislocalizated β | | 1 | | | | | |
| Without lobe(s) | | 2 | 8 | | 1 | | |
| Short α lobe | | | | | | 1 | 1 |
| Thin lobe(s) | | 1 | 1 | | | | 1 |
| Total | 40 | 41 | 31 | 19 | 46 | 28 | 23 |

**S2 Table. Phenotypes of flies that express RNAi against LpR1 or LpR2 in MB neurons under the control of the c309-Gal4 driver. Related to Fig 2.**

| | ***c309,eGFP*** | ***c309,eGFP*/+;RNAi *LpR1*/+** | ***c309,eGFP*/+;RNAi *LpR2*/+** |
|---|---|---|---|
| Normal | 26 | 11 | 20 |
| Emergence of axons from a β lobe that touch the opposite β lobe | | 7 | 3 |
| Full or partial merge of β lobes | | 2 | 1 |
| Short α lobe | | | 2 |
| Thin lobe(s) | | 1 | |
| One MB | | | 1 |
| Total | 26 | 21 | 27 |

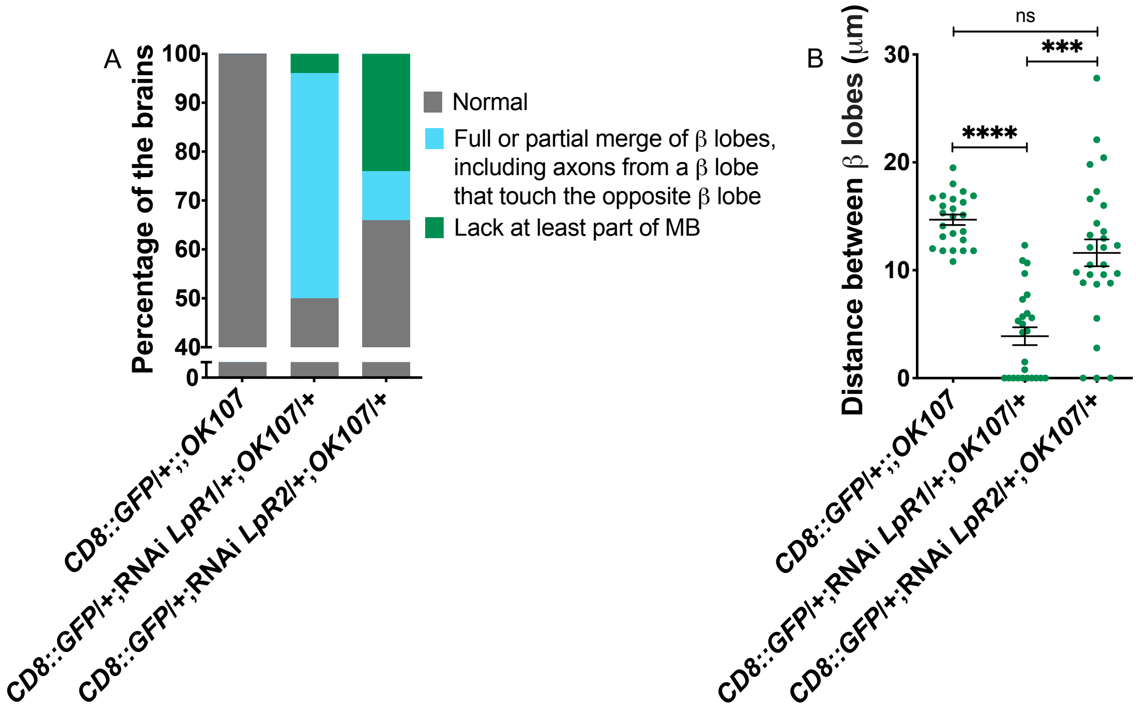


**S3 Fig. Knocking down LpR1 or LpR2 in MB neurons using the OK107 driver results in altered MB structure. Related to Fig 2.**

**S3 Table. Phenotypes of flies expressing RNAi against LpR1 or LpR2 in MB neurons under the control of OK107 driver. Related to S3 Fig.**

| | ***CD8::GFP/+;;OK107*** | ***CD8::GFP/+*;RNAi *LpR1/+;OK107/+*** | ***CD8::GFP/+*;RNAi *LpR2/+;OK107/+*** |
|---|---|---|---|
| Normal | 25 | 12 | 19 |
| Emergence of axons from a β lobe that touch the opposite β lobe | | 5 | 2 |
| Full or partial merge of β lobes | | 6 | 1 |
| Short α lobe | | 1 | |
| Thin lobe(s) | | | 4 |
| Without lobe(s) | | | 3 |
| Total | 25 | 24 | 29 |

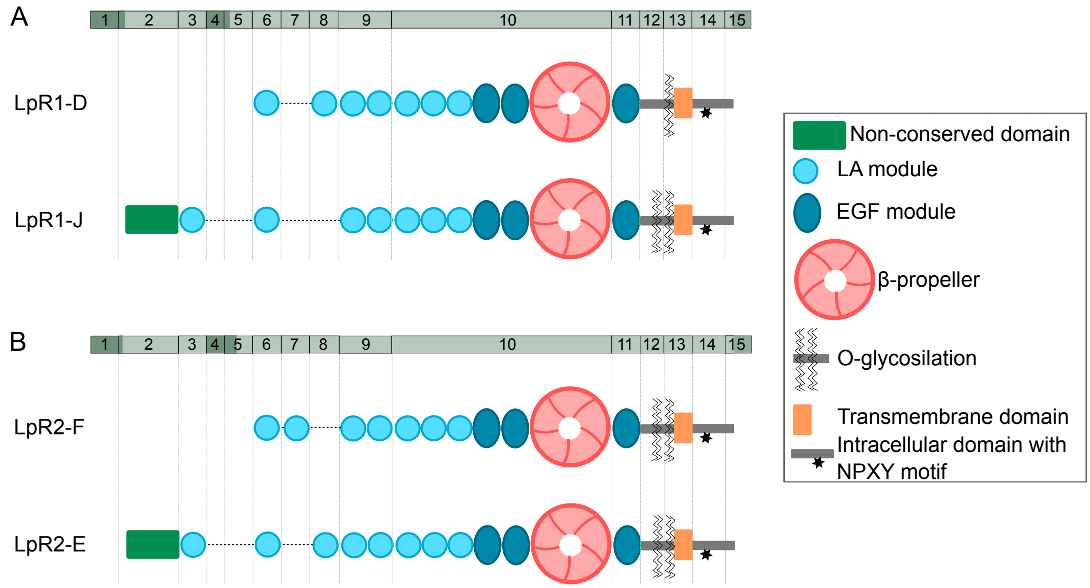


S4 Fig. Schematic representation of LpRs isoforms studied.

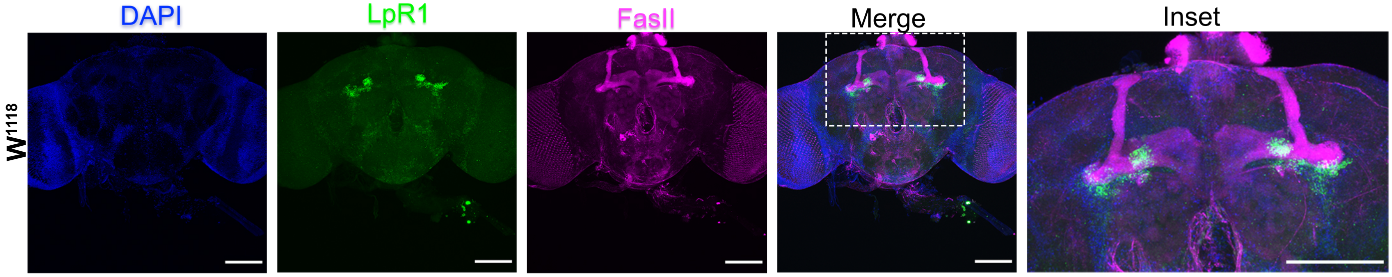


**S5 Fig. LpR1 expression in W1118 adulthood flies. Related to Fig 3.**

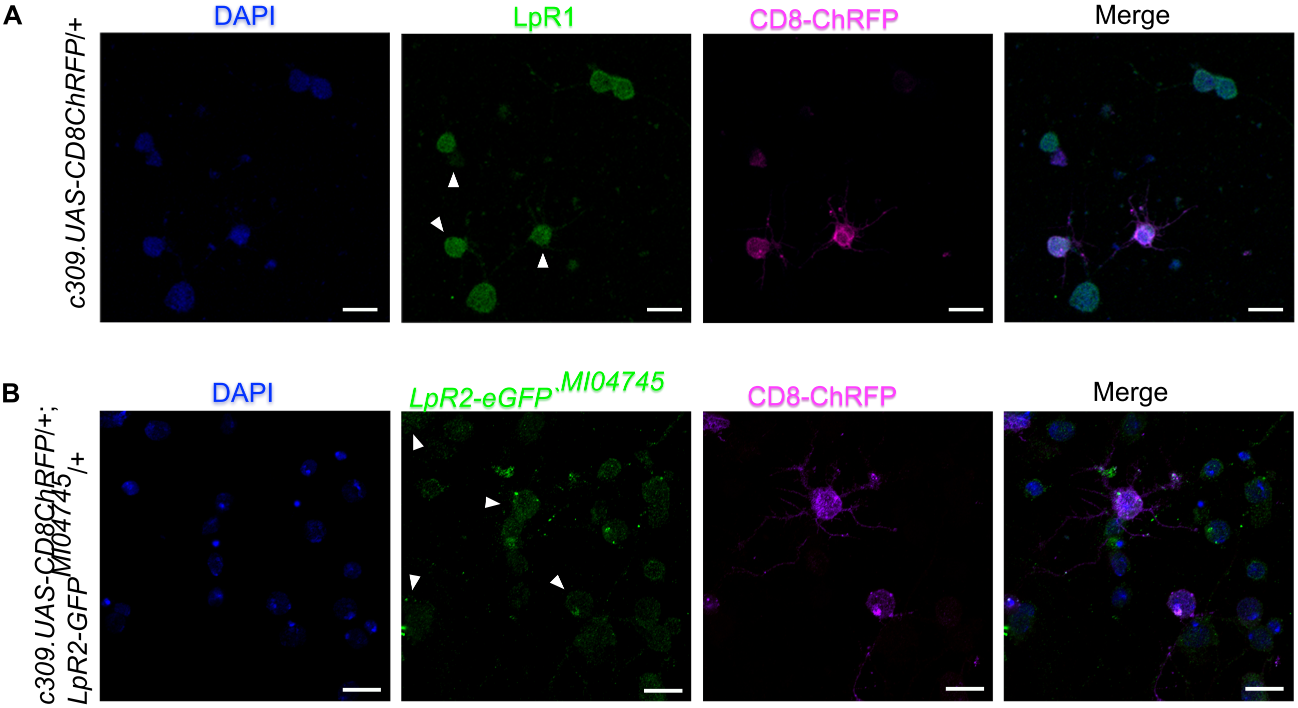


**S6 Fig. LpRs are expressed in primary culture neurons from pupal brain. Related to Fig 5.**

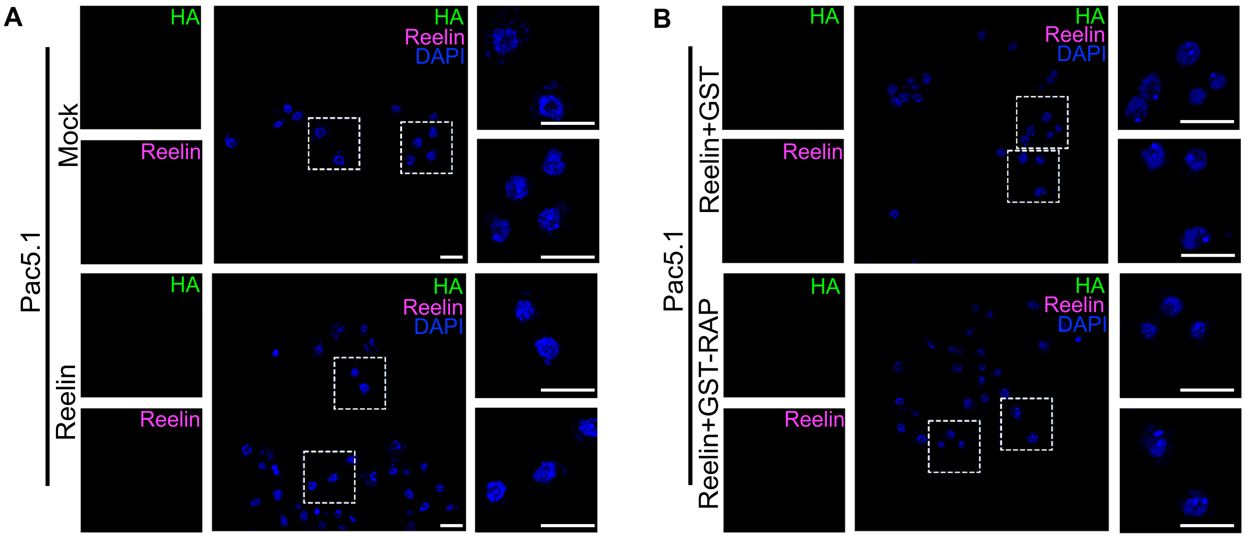


**S7 Fig. Control of the Reelin internalization. Related to Fig 6.**

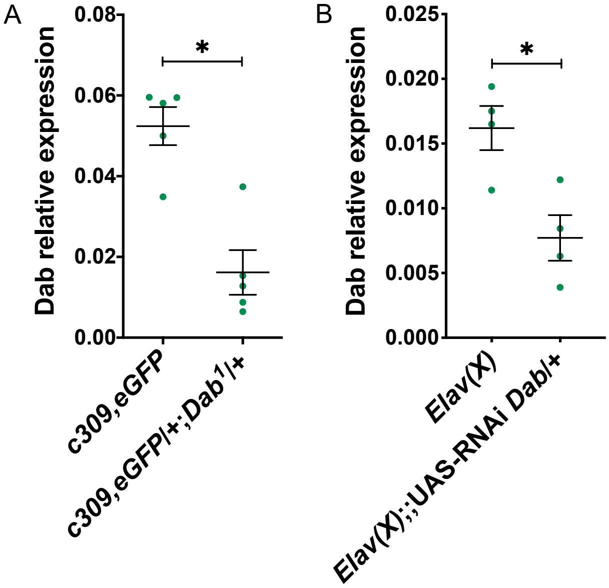


**S8 Fig. Verification of reduction in Dab expression in mutant and knockdown animals. Related to Figs 7 and 8.**

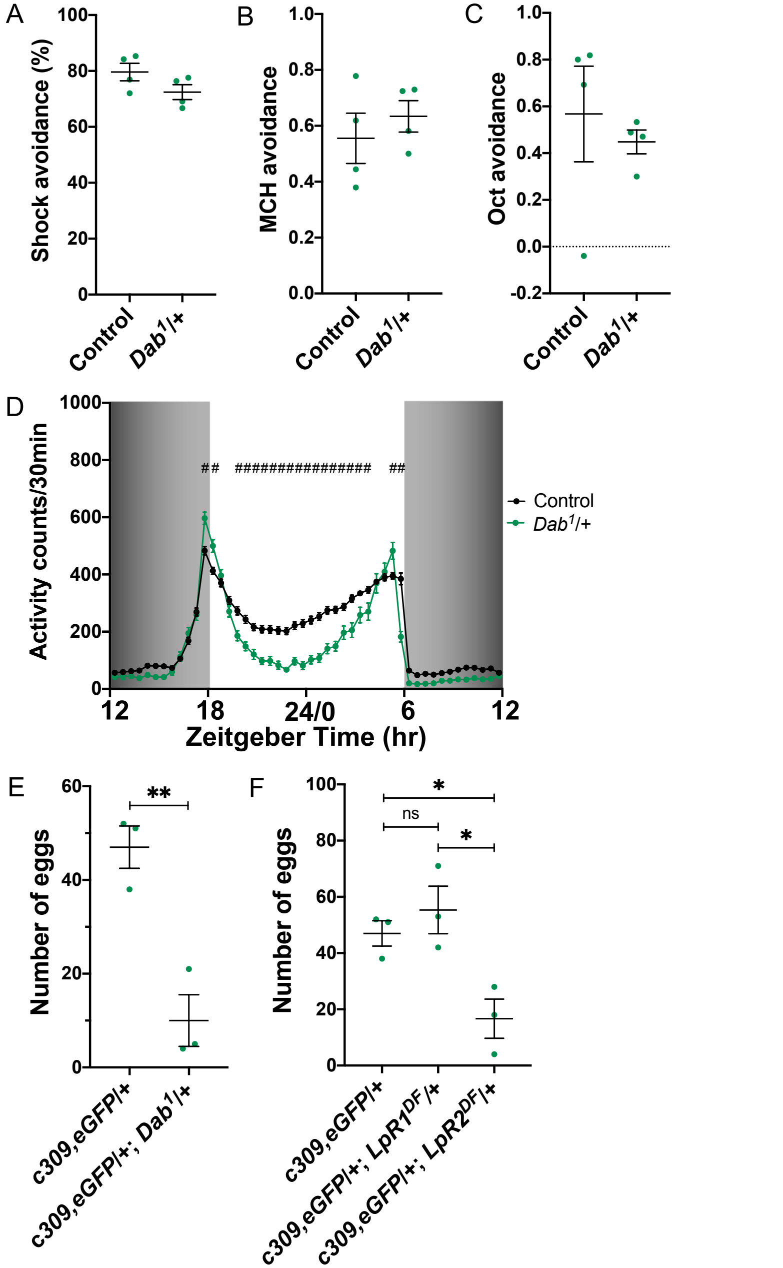


**S9 Fig. Sensory controls in Dab mutant flies. Related to Fig 8.**

**S4 Table. Phenotypes of mutant flies lacking a copy of Dab. Related to Fig 8.**

| | ***c309,eGFP*** | ***c309,eGFP*/+;*Dab*$^{1}$/+** |
|---|---|---|
| Normal | 36 | 31 |
| Mislocalizated β | | 3 |
| Split β lobes | | 1 |
| Total | 36 | 34 |

**S5 Table. Phenotypes of flies that express an RNAi against Dab in MB neurons directed by *c309-Gal4*. Related to Fig 8.**

| | ***c309,eGFP*** | ***c309,eGFP*/+;RNAi *Dab*/+** |
|---|---|---|
| Normal | 26 | 20 |
| Emergence of axons from a β lobe that touch the opposite β lobe | | 12 |
| Full or partial merge of β lobes | | 2 |
| Total | 26 | 34 |
